# Mapping heritability of obesity by brain cell types

**DOI:** 10.1101/2020.01.27.920033

**Authors:** Pascal N Timshel, Jonatan J Thompson, Tune H Pers

## Abstract

The underlying cell types mediating predisposition to obesity remain largely obscure. Here we first integrated recently published single-cell RNA-sequencing (scRNA-seq) data from >380 peripheral and nervous system cell types spanning 19 mouse organs with body mass index (BMI) genome-wide association study (GWAS) data from >450,000 individuals. Leveraging a novel strategy for integrating scRNA-seq data with GWAS data, we identified 22, exclusively neuronal, cell types from the subthalamus, midbrain, hippocampus, thalamus, cortex, pons, medulla, pallidum that were significantly enriched for BMI heritability (*P*<1.6×10^-4^). Using genes harboring coding mutations leading to syndromic forms of obesity, we replicate four midbrain cell types from the anterior pretectal nucleus, superior nucleus, periaqueductal gray and pallidum (*P*<1.7×10^-4^). Testing an additional set of 347 hypothalamic cell types, ventromedial hypothalamic steroidogenic-factor 1 (SF1) and cholecystokinin b receptor (CCKBR)-expressing neurons (*P*=4.9×10^-5^) previously implicated in energy homeostasis and glucose control and three cell types from the preoptic area of the hypothalamus and the lateral hypothalamus enriched for BMI GWAS associations (P<4.9×10^-5^). Together, our results suggest brain nuclei regulating integration of sensory stimuli, learning and memory are likely to play a key role in obesity and provide testable hypotheses for mechanistic follow-up studies.

## INTRODUCTION

Identification of genes and cell types underlying susceptibility to human obesity remains a critically important step towards a better understanding of mechanisms causing the disease ^1^. Studies of monogenic obesity syndromes and rodent models of obesity have identified melanocortin signaling circuits in the mediobasal and paraventricular hypothalamus as key components in energy homeostasis and obesity ^2–4^. Yet growing evidence suggests that susceptibility to obesity is distributed across numerous brain areas that receive signals emanating from internal sources (*e.g.,* viscerosensory input from the gastrointestinal tract) or external stimuli (*e.g.,* the sight or smell of food) that act in concert to regulate feeding behavior and energy stores ^5–7^. However, despite an increasing number of genes, cell types and neuronal circuits being implicated in murine energy homeostasis, the identity of brain cell types that drive susceptibility to human obesity remains largely unknown and a systematic assessment of cell types’ relevance in obesity is currently lacking.

In recent years, genome-wide association studies (GWAS) have identified about a thousand common (minor allele frequency, MAF≥0.1) single nucleotide polymorphisms (SNPs) that associate with body mass index (BMI, defined as weight in kilogram divided by height in meters squared), a heritable and commonly used proxy phenotype for obesity ^8, 9^. In general, the far majority of trait-associated SNPs are located in regulatory regions and hence, unlike coding variants, tagging genetic intervals (or *loci*) rather than implicating specific genes. Importantly, these loci represent an unbiased set of biological sign posts to genes and biological mechanisms underlying susceptibility to obesity ^10^.

Genetic variants with rare frequencies (MAF<0.1) that are typically too low to be captured in GWAS are thought to contribute ∼50% to the heritability of BMI ^11^. Many such variants are coding mutations ^11^ and hence well-suited to identify causal genes underlying obesity. Lately, rare variant association studies have identified 14 coding variants across 13 genes in an exome chip analysis across >750,000 individuals ^12^. Interestingly, these genes, with the exception of *MC4R, KSR2* and *GIPR,* have not previously been implicated in obesity, suggesting that key biologic mechanisms underlying obesity have yet to be identified.

Given that a majority of obesity-associated gene variants likely regulate gene expression rather than impact protein function, gene expression data provide an effective scaffold to inform GWAS data for obesity and other traits ^13–16^. In 2016, we used microarray-based gene expression data to show that genes in BMI GWAS loci are predominantly expressed in the brain ^8^ and we recently leveraged mouse single-cell RNA-sequencing (scRNA-seq) to implicate mediobasal hypothalamic cell types in obesity ^17^. The growing number of BMI GWAS loci and genes implicated through rare-variant association studies of common and syndromic forms of obesity, in conjunction with the growing number of large-scale scRNA-seq atlases, provide a unique opportunity to systematically uncover genes and cell types underlying biological circuits regulating susceptibility to human obesity.

Here we developed two computational toolkits for human genetics-driven identification of cell types underlying disease and leveraged them to systematically identify cell types enriching for obesity susceptibility by combining publicly available BMI GWAS summary statistics from >450,000 individuals with scRNA-seq data spanning 380 cell types representing adult mouse organs especially the nervous system and 347 cell types from the adult mouse hypothalamus.

## RESULTS

### Devising a robust cell type expression specificity metric and prioritization framework

Similar to previous approaches ^17–20^, we hypothesized that cell types exhibiting detectable expression of genes colocalizing with BMI GWAS loci are more likely to underlie obesity than cell types in which the expression of genes is low. Based on this reasoning, we developed CELLECT (**Cell** type **E**xpression-specific integration for **C**omplex **T**raits) and CELLEX (**Cell** type **EX**pression-specificity), two toolkits for genetic identification of likely etiologic cell types. Given GWAS summary statistics and scRNA-seq data, CELLECT can quantify the enrichment of heritability in or near genes specifically expressed in a given cell type using established genetic prioritization models, such as S-LDSC ^13^, RolyPoly ^14^, DEPICT ^15^ or MAGMA covariate analysis ^18^ (Methods; **Fig. 1a**). Importantly, whereas previous frameworks for genetic prioritization of cell types have either relied on non-polygenic models ^17^, used binary or discrete representations of cell type expression ^13, 18^ or used average expression profiles ^20^, CELLECT uses a robust continuous representation of cell type expression. In **Supplementary Notes 1,2** we provide a detailed discussion of our model, its assumptions and relationship to the ‘omnigenic’ model hypothesis ^21, 22^. Conjointly, CELLEX was built on the observation that different measures of gene expression specificity (ES) provide complementary information and therefore combines four ES metrics (see Methods) into a single measure (*ES_μ_*) representing the score that a gene is specifically expressed in the given cell type (Methods; **Fig. 1b**). Our ES approach was tested and validated using the Tabula Muris dataset ^23^, a Smart-Seq2 scRNA-seq dataset derived from 20 organs from adult male and female mice, and the Mouse Nervous System dataset ^24^, derived from 19 central and peripheral nervous system regions from late-postnatal male and female mice. Based on a total of 53,760 cells, we computed gene expression specificity for the four metrics and combined them into *ES_μ_* across four cell types with known marker genes and found that *ES_μ_* correctly identified them as being among the most specifically expressed genes (**Fig. 1d,e**; **Supplementary Results**). We identified a median of 2,934 specifically expressed genes per cell type and hierarchical clustering of cell types based on the *ES_μ_* estimates largely reproduced the cell type dendrograms from the respective original publications ^23, 24^, confirming that our ES approach enables cell types profiles to be compared across studies and technologies (**Supplementary Fig. 1,2**). In **Supplementary Note 3** we provide a detailed description of the CELLEX workflow, its assumptions and we use re-sampling to demonstrate the robustness of *ES_μ_* compared to individual ES metrics. Both CELLECT and CELLEX have been implemented and released as open-source packages for Python programming languages (URLs). Here, we used CELLECT with S-LDSC as the genetic prioritization model to quantify the effects of cell type ES on BMI heritability. For each cell type, we reported the *P*-value for the one-tailed test for positive contribution of the cell type ES to trait heritability (conditional on a ‘baseline model’ that accounted for the non-random distribution of heritability across the genome, see Methods).

**Fig. 1.**
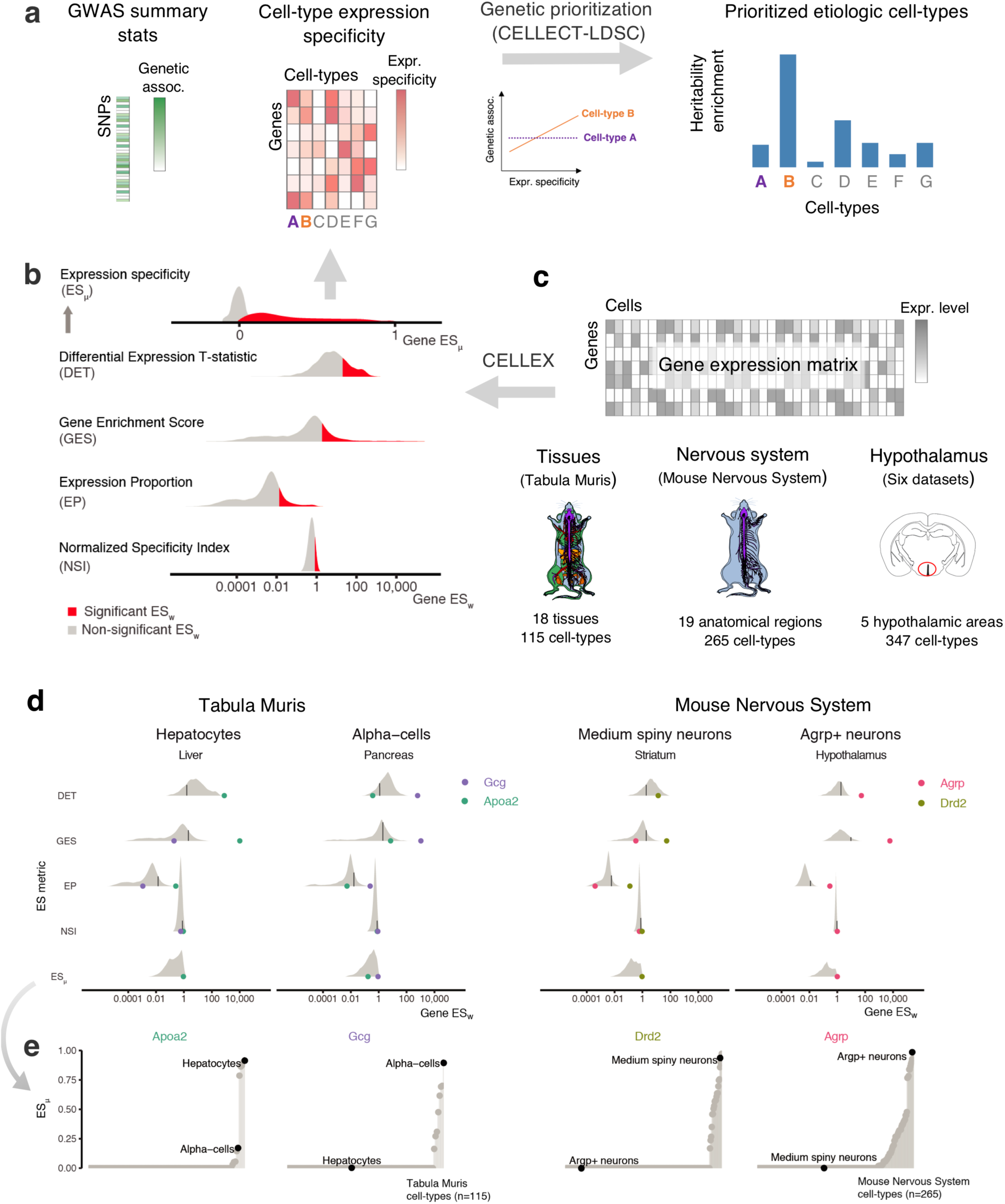
Overview of CELLECT and CELLEX and main datasets used. **a**, CELLECT quantifies the association between common polygenetic GWAS signal (heritability) and cell type expression specificity (ES) to prioritize relevant etiological cell types. As input to CELLECT, we used BMI GWAS summary statistics derived from analysis of UK Biobank data (N>457,000 individuals) and ES calculated using CELLEX. **b,** CELLEX uses a ‘wisdom of the crowd’ approach by averaging multiple ES metrics into *ES_μ_*, a robust ES measure that captures multiple aspects of expression specificity. Prior to averaging ES metrics, CELLEX determines the significance of individual ES metric estimates (*ES_w_*), indicated by the red and gray colored areas. **c**, scRNA-seq datasets analyzed in this study. In total the associations between 727 cell types and BMI heritability were analyzed. Anatograms modified from gganatogram ^70^. **d**, Example of the CELLEX approach for selected cell types and relevant marker genes. The log-scale distribution plot of *ES_w_* illustrate differences of ES metrics. For each ES metric distribution, a black line is shown to indicate the cut-off value for *ES_w_* significance. In most cases, the ES metrics identified the relevant marker gene as having a significant *ES_w_*. In all cases, the marker gene is correctly estimated as having *ES_μ_*∼1. We note that the majority of genes have *ES_μ_*=0 and were omitted from the log-scale plot. **e**, *ES_μ_* plots showing the specificity and sensitivity of our approach. The plots depict *ES_μ_* for the genes shown in panel (d) across all cell types in the respective datasets. For each marker gene the relevant cell type has the highest *ES_μ_* estimate (high sensitivity) and cell types in which the given gene is likely to have a lesser role have near zero *ES_μ_* estimates (high specificity). BMI, body mass index; GWAS, genome-wide association study; UK, United Kingdom; scRNA-seq, single-cell RNA-sequencing.

### BMI variants enrich for central nervous system rather than peripheral cell types

Using BMI GWAS summary statistics from a GWAS analysis of the UK Biobank ^25^ comprising >457,000 individuals ^26^ and the Tabula Muris cell types, we first assessed whether we could replicate the exclusive enrichment of BMI GWAS variants in brain tissues as reported by Locke *et al*. ^8^ **(Supplementary Tables 1-3)**. Applying CELLECT to the 115 – mostly peripheral – cell types, we identified two significantly enriched cell types, namely neurons and oligodendrocyte precursor cells (Bonferroni correction-based false-discovery rate, FDR<0.05; **Fig. 2a, Supplementary Table 4).** When rerunning CELLECT conditioning the neuron cell type, the oligodendrocyte precursor cell type was not significant any longer, emphasizing that we primarily observed a neuron signal for GWAS BMI variants. In order to verify that our approach, in general, could identify relevant cell types for complex traits, we computed enrichments for nine GWAS including cognitive, psychiatric, neurological, immunological, lipid and anthropometric traits and disorders, and found that CELLECT prioritized etiologically relevant cell types across all six categories (**Fig. 2b, Supplementary Table 4**). Thus, cortical neurons were prioritized for cognitive traits and psychiatric disorders (educational attainment, intelligence, schizophrenia), neuronal cell types for insomnia, immune cells for multiple sclerosis and rheumatoid arthritis, growth-related cell types for waist-to-hip ratio (adjusted for BMI) and height, and hepatocytes for low-density lipoprotein levels (see **Supplementary Table 4** for results across additional 29 traits). These data establish the ability of this approach to validate previous evidence ^8^ that BMI variants tend to colocalize with genes specifically expressed in neurons, while also demonstrating that CELLECT is able to prioritize relevant cell types across a number of complex traits.

**Fig. 2.**
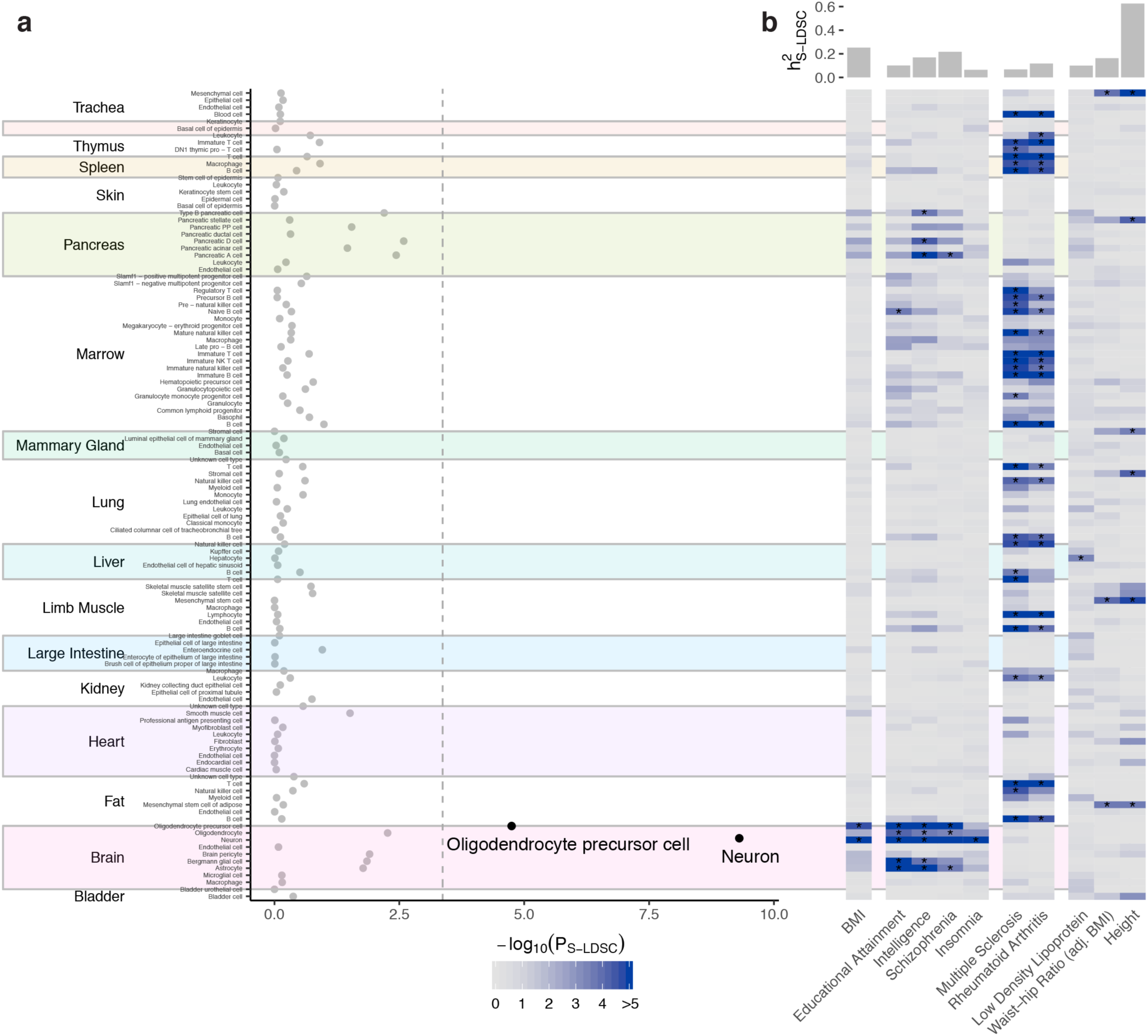
Cell type prioritization across 19 tissues highlights the key role of the brain in obesity. **a**, Prioritization of 115 Tabula Muris cell types identified two cell types from the brain as significantly associated with BMI, namely oligodendrocyte precursor cells and neurons (shown in black; Bonferroni significance threshold, *P*_S-LDSC_<0.05/115). **b**, Heatmap of cell type prioritization for multiple GWAS traits. BMI results (first column) are the same as in panel (a) projected onto the heatmap. The four brain-related traits (second column) were associated with cell types in the brain, the two immune traits (third column) were associated with immune cells, and anthropometric traits (fourth column) were associated with mesenchymal stem cells, which are progenitor cells for muscle, bone and fat. Asterisks (*) mark cell types passing the per-trait Bonferroni significance threshold. The top bar plot shows the estimated trait heritability. S-LDSC, stratified-linkage disequilibrium score regression; h^2^_S-LDSC_, trait SNP-heritability.

### A distributed set of neuronal cell types enrich for obesity susceptibility

We next assessed whether we could identify specific CNS cell types enriching for BMI-associated variants. Applying CELLEX and CELLECT on 265 cell types from the mouse nervous system, we identified 22 enriched cell types annotated to eight brain regions (**Fig. 3a; Supplementary Table 5,6**). To assess the specificity of the BMI signal in these 22 cell types, we computed enrichments for the panel of nine other well-powered traits. As expected, none of the five traits primarily caused by peripheral etiologies enriched for any nervous system cell type and several of 22 BMI GWAS-enriched cell types also enriched for cognitive traits and psychiatric disorders (**Fig. 3b; Supplementary Table 5**). Sixteen of the 22 cell types were also enriched “intelligence” and “worry”, two traits genetically anticorrelated with obesity (overlapping sets of associated loci with opposite effect sizes) ^27, 28^ (**Supplementary Table 7**).

**Fig. 3.**
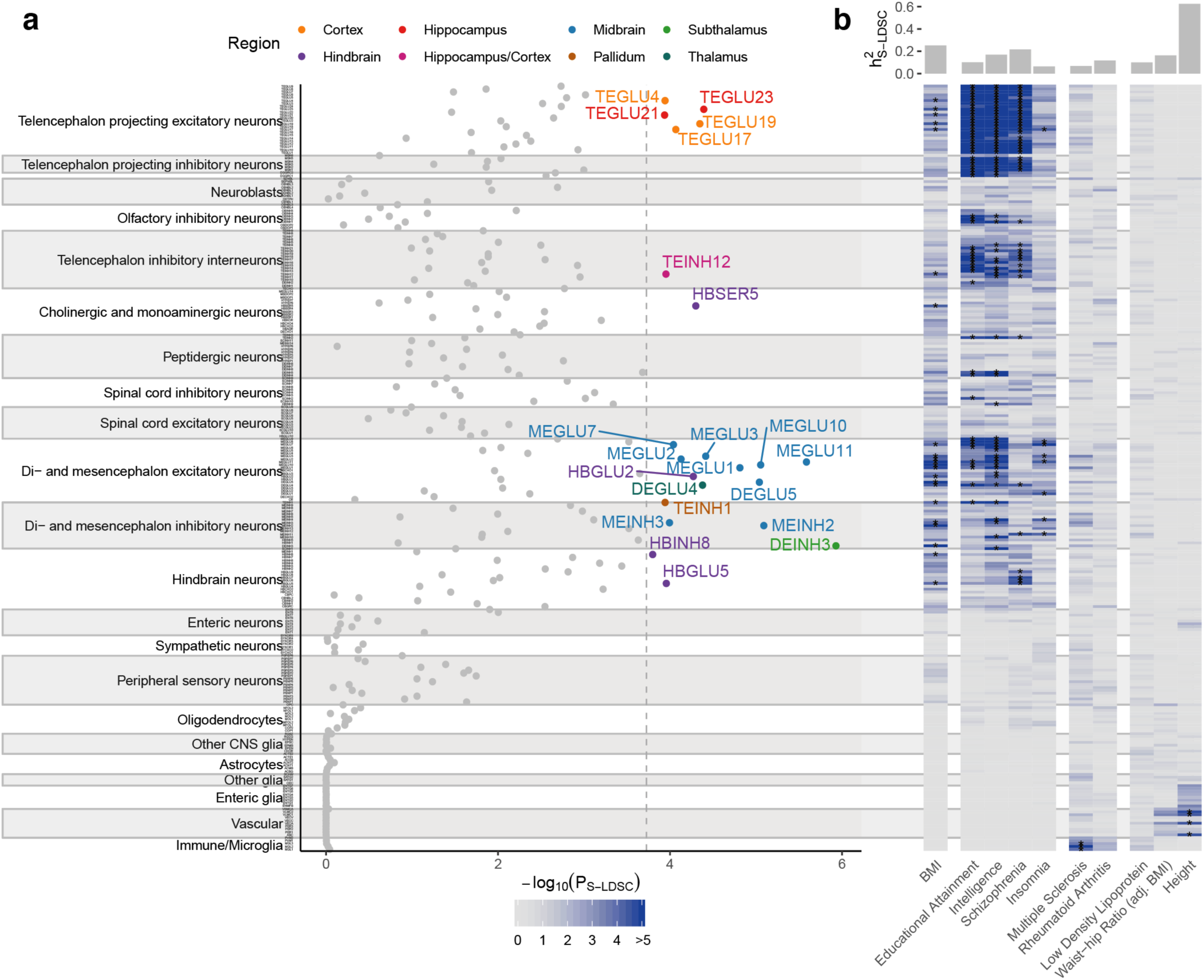
Cell type prioritization of mouse nervous system cell types highlights cell types outside canonical energy homeostasis circuits. **a**, Prioritization of 265 mouse nervous system cell types identified 22 cell types from eight distinct brain regions as significantly associated with BMI (**Supplementary Table 5** for details). The highlighted cell types passed the Bonferroni significance threshold, *P*_S-LDSC_<0.05/265). Cell types were grouped by the taxonomy described in Zeisel et al. (2018). **b**, Heatmap of cell type prioritization for multiple GWAS traits. The four brain-related traits (second column) were primarily associated with cortical neurons (telencephalon projecting and interneuron cell types) and did not overlap with the BMI-associated cell types. The two immune traits (third column) were associated with microglia, and anthropometric traits (fourth column) were predominantly associated with vascular cell types. Asterisks (*) mark cell types passing the per-trait Bonferroni significance threshold. The top bar plot shows the estimated trait heritability.

Similar to previous work, we did not find any enrichment of genetic variants associated with BMI in non-neuronal cell types ^17, 20^ nor did we detect enrichment for a particular type of neurotransmitter type (**Supplementary Fig. 5)**. Weighted correlation network analysis (WGCNA, ^29^) on expression data from each of the 22 BMI-enriched cell types identified no significant modules (**Supplementary Fig. 4; Supplementary Tables 8,9;** top associated module, *P*=1.88×10^-4^; FDR≤0.1). These findings emphasize that the BMI-associated variants most likely are distributed across hundreds of genes rather than the relatively limited number of genes captured in cell type-specific WGCNA modules (see **Supplementary Note 4** for a discussion on limitations of identifying gene co-expression networks from cell type scRNA-seq data).

To assess the dependence of the results on a given enrichment methodology and BMI GWAS, we re-computed enrichments using the Yengo *et al.* ^9^ and Locke *et al.* ^8^ BMI GWAS summary statistics and the MAGMA tool ^30^ (**Supplementary Tables 5,10**). We observed that the results were robust to different GWAS sample sizes and inclusion of Metabochip arrays (Yengo *et al.* and Locke *et al.* GWAS Pearson’s *R*=0.98 *and R*=0.83, respectively), and largely invariant to the enrichment methodology used (Pearson’s *R*=0.82; **Supplementary Fig. 5**). Finally, during finalizing this work another study focused on Parkinson’s disease, reported BMI GWAS enrichments for the same mouse nervous system cell types (overlap; 6/10) ^19^. Together, these results demonstrate that BMI-associated variants are likely to exert their effect across multiple, exclusively neuronal cell types, several of which enrich for cognitive traits and psychiatric disorders genetically correlated with obesity.

### The enriched neuronal cell types share transcriptional similarities

The 22 cell types mapped to eight brain regions, namely the subthalamus, midbrain, hippocampus, thalamus, cortex, pons, medulla and pallidum (**Fig. 4a**). We next assessed whether shared transcriptional signatures could explain the enrichments across the 22 cell types. Towards that aim, we first clustered all cell types based on their genes’ *ES_μ_* values and, expectedly, found that midbrain cell types overall grouped by their neuroanatomical proximities and neurotransmitter types into midbrain, hindbrain, hippocampus/cortex clusters (**Fig. 4b**). A notable exception was the DEINH3 cell type (isolated from the hypothalamus region and subsequently remapped to the subthalamic nucleus by Zeisel *et al.*) which grouped with the midbrain cell types. To further assess the transcriptional similarity between the enriched cell types, we computed enrichments conditioned on each prioritized cell type individually (Methods). Contrary to our expectations, we found that none of the other cell types remained significant when conditioning on the top-ranked subthalamic cell type DEINH3 (**Supplementary Fig. 6,7; Supplementary Table 11).** Together these results indicate the brain cell types enriching for BMI GWAS signal, despite their neuroanatomical differences, share transcriptional signatures related to obesity, which current methods are not able to disentangle.

**Fig. 4.**
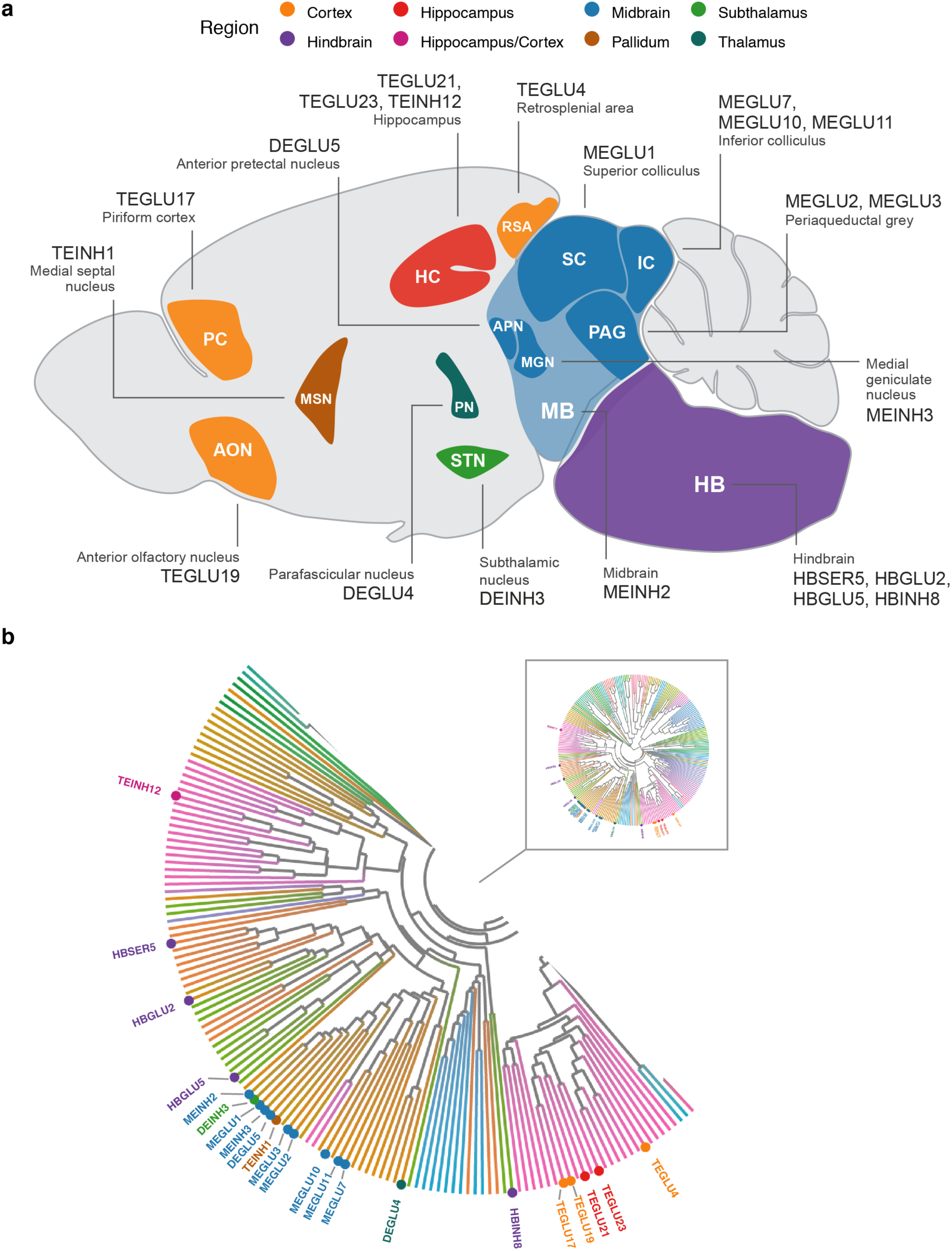
Neuroanatomical location and transcriptional similarity of brain cell types enriching for BMI GWAS variants. **a,** Sagittal mouse brain view showing the 22 enriched cell types. The first two letters in each cell type label denote the developmental compartment (ME, mesencephalon; DE, diencephalon; TE, telencephalon), letters three to five denote the neurotransmitter type (INH, inhibitory; GLU, glutamatergic) and the numerical suffix represents an arbitrary number assigned to the given cell type. **b**, Circular dendrogram showing the similarity of all Mouse Nervous System dataset cell type expression specificity (*ES_μ_*) values. Dendrogram edges colored by taxonomy described in Zeisel et al. (2018). Expectedly, the cell types clustered according to their neuroanatomical origin. For clarity only the labels of the 22 BMI GWAS enriched cell types are shown.

### Ventromedial hypothalamic Sf1- and Cckbr-expressing cells enrich for BMI GWAS

The total number of cell types in the hypothalamus has been significantly underestimated ^31^, therefore to assess whether the lack of enrichment for hypothalamic cell types was due to sparse sampling of hypothalamic cells in the Mouse Nervous System dataset, we computed enrichments for an additional set of 347 cell types sampled from the mediobasal hypothalamus ^17^, the ventromedial hypothalamus ^31^, the lateral hypothalamus ^32^, the preoptic nucleus of the hypothalamus ^33^ and the entire hypothalamus ^34, 35^ (**Supplementary Table 12**). We identified four non-overlapping significantly enriched cell types, namely a ventromedial hypothalamic glutamatergic cell type (ARCME−NEURO29; *P=*4.9×10^-5^) expressing Sf1 (ES*_μ_*=0.98 and ES*_μ_*=0.99), Cckbr (cholecystokinin B receptor; ES*_μ_*=0.98, ES*_μ_*=0.95); a glutamatergic cell type from the lateral hypothalamus (LHA-NEURO20; *P*=4.9×10^-5^); two cell types from the preoptic area of the hypothalamus (POA-NEURO21 and POA-NEURO66; *P<*1.0×10^-4^; **Fig. 5**; **Supplementary Table 13,14**). Interestingly, ventromedial hypothalamic neurons have previously been implicated in control of both body fat mass and blood glucose levels; disrupted leptin signaling in Sf1-expressing ventromedial hypothalamic neurons renders mice more susceptible to diet-induced weight gain ^36^ and activation of ventromedial hypothalamic Sf1 neurons causes hyperglycemia ^37^. The two cell types also expressed Bdnf (ES*_μ_*=0.91, ES*_μ_*=0.99); mutations in *BDNF* and its receptor is a known cause of monogenic obesity in humans and, in mice, Bdnf signaling is required for normal energy homeostasis and glucoregulatory control ^38^. (Bdnf and Ntrk2 were also specifically expressed in TEINH12 cell type, a cholecystokinin (Cck)-expressing interneuron, enriched in the Mouse Nervous System analysis, see **Supplementary Results**.) Notably, there was no significant enrichment for Agrp and Pomc neurons (coding mutations in *POMC* can cause syndromic obesity in humans), however, two Pomc cell populations were nominally enriched (HYPR-NEURO24 ^35^, *P*=0.002; ARCME-NEURO21 ^17^, *P*=0.01). In sum, although we identified enrichment in four – exclusively neuronal cell types – the overall BMI GWAS enrichment in hypothalamic cell types was surprisingly limited.

**Fig. 5.**
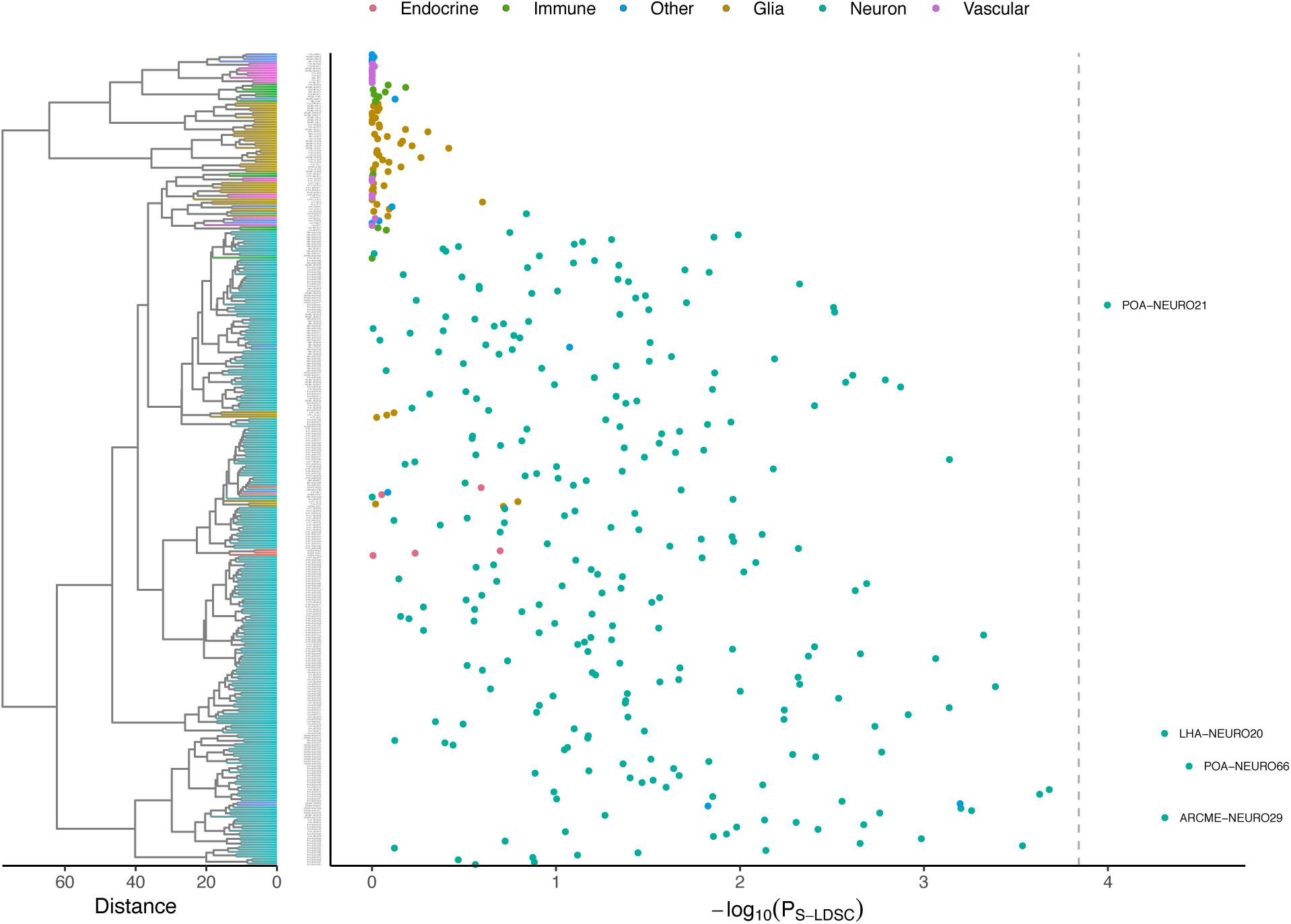
Hypothalamic cell type enriching for BMI GWAS variants. **a**, BMI GWAS enrichments across 347 hypothalamic cell types derived from studies the arcuate nucleus and median eminence complex, the ventromedial hypothalamus, the lateral hypothalamus, the preoptic nucleus of the hypothalamus and the entire hypothalamus. The dendrogram (left) shows the similarity of all hypothalamic cell types (Ward’s hierarchical clustering on *ES_μ_* values); neuronal and non-neuronal cell types clustered together across studies. For each study, CELLEX and CELLECT were run individually, and subsequently all cell types were pooled and significance was determine based on Bonferroni correction (*P*<0.05/347). Four cell types were significantly enriched, namely POA-NEURO66 (top marker Reln in ref. ^33^) and POA-NEURO21 (top markers Cck and Ebf3 in ref. ^33^) from the preoptic area of the hypothalamus, ARCME-NEURO29 (top markers Sf1 and Adcyap1 in ref. ^17^) from the arcuate nucleus and median eminence complex, and LHA-NEURO20 (top marker genes Ebf3 and Otb in ref. ^32^) from the lateral hypothalamus. POA, preoptic area of the hypothalamus; LHA, lateral hypothalamus; ARCME, arcuate nucleus and median eminence complex; S-LDSC, stratified-linkage disequilibrium score regression.

### Genes with known links to human obesity genes implicate the dorsal midbrain

As genes that have been associated with obesity through sequencing and large-scale exome chip association analysis have been identified independently of the results from GWAS, we reasoned that we could use them to validate the cell types enriched for BMI GWAS signal. Combining a set of 39 obesity genes associated with syndromic forms of obesity (obtained from Turcot *et al.* ^12^) and 13 genes with rare but high-effect size mutations implicated by the Turcot *et al.* study, we computed the enrichment of a total set of 50 unique obesity genes (**Supplementary Table 15**) within all 265 Mouse Nervous System cell types. We identified 15 significantly enriched cell types (one-sided Mann-Whitney U test, FDR<0.05) of which four replicated cell types from the BMI GWAS analysis (DEGLU5, MEGLU1, MEGLU2 and TEINH1 from the anterior pretectal nucleus, the superior colliculus, the periaqueductal grey and the pallidum; *P*<1.7×10^-4^; (**Supplementary Fig. 8, Supplementary Table 16**). Among the remaining 11 cell types, 10 originated from areas implicated by the CELLECT analysis, namely the midbrain (MBDOP1, periaqueductal grey; MEINH4, the anterior pretectal nucleus; MEGLU6, superior colliculus; MEGLU14, the dorsal raphe nucleus), the hypothalamus (HYPEP1, lateral hypothalamus; DEINH5, the medial preoptic nucleus and bed nuclei of the stria terminalis anterior division; and the dorsomedial nucleus of the hypothalamus), the medulla (HBSER4, nucleus raphe medulla; HBNOR, A1-2 noradrenergic cell groups) and the thalamus (DEINH4, paraventricular nucleus of the thalamus; **Fig. 6a**). We observed a significant correlation between the rare variant obesity geneset and CELLECT cell type enrichment results (Pearson‘s *R*=0.58*, P*=1.7×10^-25^; **Supplementary Fig. 9a**), which further underscored the validity of our findings and, as previously reported ^8^, that genes implicated in syndromic obesity tend to colocalize with BMI-associated GWAS loci (**Supplementary Fig. 9b**).

**Fig. 6.**
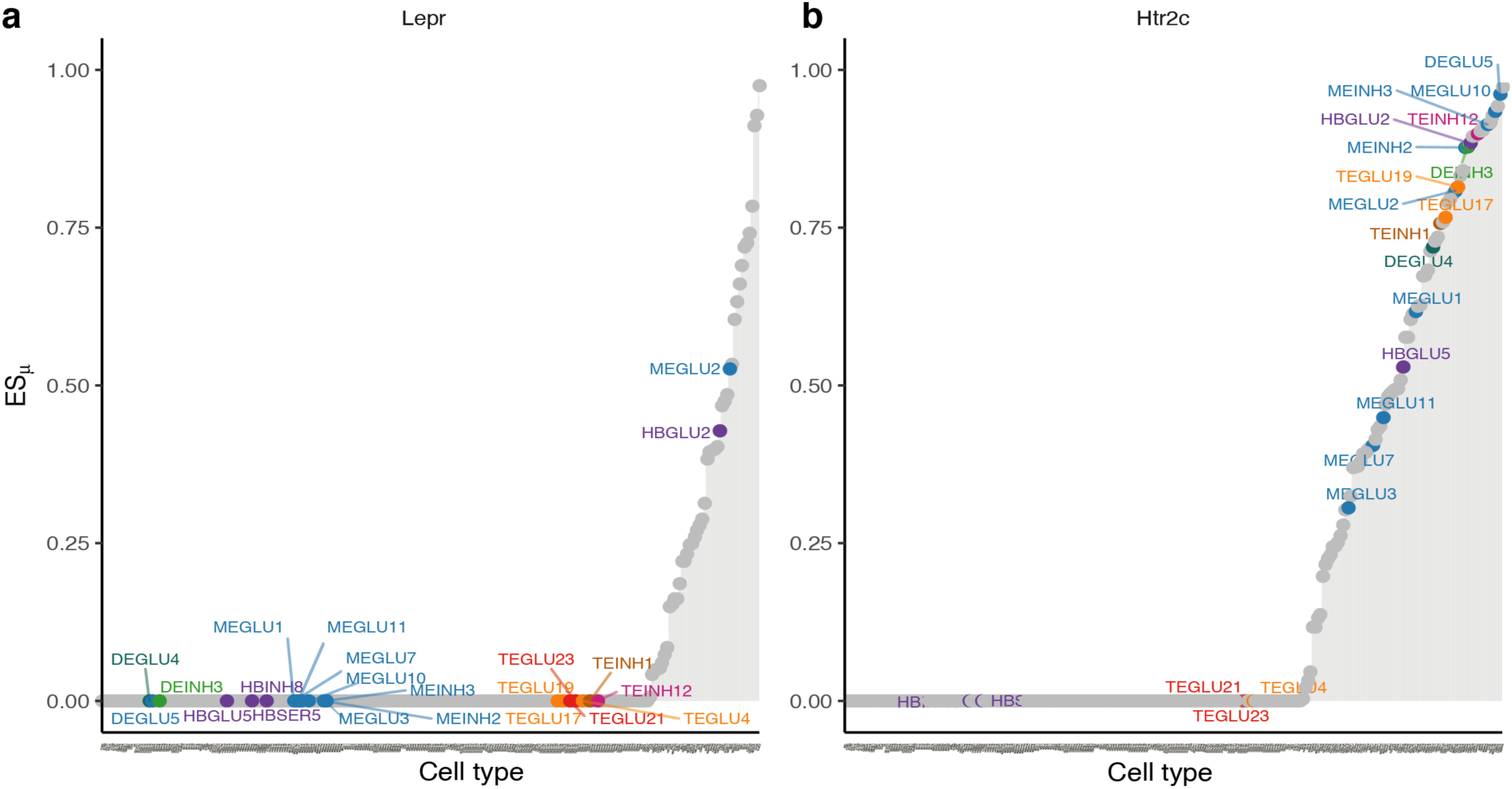
Expression specificity of the leptin- and serotonin receptors across BMI GWAS enriched cell types. **a**, In the lipostatic model of obesity (originally defined by Kennedy (1953), circulating concentrations of the leptin hormone signal the amount of energy stored in fat cells to the brain. The plot shows gene *ES_μ_* (y-axis) for each cell type (x-axis, ordered by increasing values of expression specificity, *ES_μ_*) with BMI-prioritized cell types highlighted. In our analysis, only two of the 22 BMI GWAS enriched cell types specifically expressed the leptin receptor (MEGLU2, periaqueductal grey; and HBGLU2, nucleus of the solitary tract). **b**, Seventeen of the 22 BMI GWAS enriched cell types specifically expressed the serotonin (5-htr2c) receptor. The strongest enrichment was observed for DEGLU5, a glutamatergic cell type from the anterior pretectal nucleus. *ES_μ_*, expression specificity.

Interestingly, the leptin receptor, which regulates key energy homeostatic processes in the hypothalamus and when defective may cause syndromic obesity ^39^ was only specifically expressed in two out of the 22 BMI GWAS-enriched cell types, namely glutamatergic cells from the periaqueductal grey and anterior nucleus of the solitary tract (**Fig. 6a**). By contrast, 17 of the enriched cell types expressed the serotonin receptor 5-Htr2c (5-hydroxytryptamine receptor 2C), a known regulator of energy and glucose homeostasis ^40^ and the target for current obesity pharmacotherapy ^41^ (**Fig. 6b**). 5-Htr2c was most specifically expressed in the anterior pretectal nucleus (DEGLU5, ES*_μ_*=0.96), the cell type among our results which most specifically expressed Pomc (ES*_μ_*=0.41). Mice lacking the Htr2c in Pomc neurons are resistant to 5-Htr2c agonist Lorcaserin-induced weight loss ^40^ (For ES_μ_ plot of other selected genes, please refer to **Supplementary Fig. 10**).

Together these results indicate that susceptibility to obesity conferred by common variants, while enriching for some hypothalamic cell types such as VMH Sf1-expressing neurons, is distributed across a mosaic of neuronal cell types of which numerous are involved in regulating integration of sensory stimuli, learning and memory.

## DISCUSSION

Here, we developed two scRNA-seq computational toolkits called CELLEX and CELLECT and applied them to scRNA-seq data from a total of 727 mouse cell types, from mostly adult mice, to derive an unbiased map of cell types enriching for human genetic variants associated with obesity. In total, we identified 26 neuronal cell types, which, in line with previous considerations ^5^, demonstrate that susceptibility to human obesity is likely to be distributed across multiple, mainly neuronal, cell types across the brain, rather than being mostly restricted to a limited number of canonical energy homeostasis- and reward-related brain areas in the hypothalamus, midbrain and hindbrain. Among the enriched hypothalamic cell types, we identified VMH Sf1- and Cckbr-expressing neurons, which previously have been implicated in glucose and energy homeostasis.

### Processing of sensory stimuli and feeding behavior

Several of the BMI GWAS enriched cell types localized to nuclei integrating sensory input and directed behavior. The inferior colliculus (implicated by MEGLU7, MEGLU10 and MEGLU11) and medial geniculate nucleus (MEINH3) process auditory input, the superior colliculus for translation of visual input into directed behavior (MEGLU1 and MEGLU6), the anterior pretectal nucleus processes somatosensory input (DEGLU5 and MEINH4), and the piriform cortex (TEGLU17) and anterior olfactory nucleus (TEGLU19) processing odor perception. The superior colliculus and anterior pretectal nucleus have both implicated in predatory behavior ^42, 43^ and project to the zona incerta, a less well-described brain area situated between the thalamus and hypothalamus that receives direct input from mediobasal hypothalamic Pomc neurons ^44^. In rats, lesioning of the zona incerta impairs feeding responses ^45^ while, conversely, in mouse models, optogenetic stimulation of GABAergic neurons in the zona incerta leads to rapid, binge-like eating and body weight gain ^46^. Activation of projections from the hypothalamic preoptic nucleus (DEINH5; POA-NEURO21, POA-NEURO66) to the ventral periaqueductal gray (MEGLU2 and MBDOP1) induce object craving ^47^, whereas pharmacological inactivation of the periaqueductal gray decreases food consumption ^48^. Together, these findings suggest that susceptibility to obesity is enriched in cell types processing sensory stimuli and directing actions related to feeding behavior and opportunity.

### Evidence supporting a key role of the learning and memory in obesity

Feeding is not an unconditioned response to an energy deficiency but rather reflecting behavior conditioned by learning and experience ^49^. We previously showed that genes in BMI GWAS loci enrich for genes specifically expressed in hippocampal post-mortem gene expression data ^8^. In this work, we identified specific brain cell types supporting a role of memory in obesity. First, the parafascicular nucleus (DEGLU4), when lesioned in mice, reduces object recognition memory ^50^. Second, the retrosplenial cortex (TEGLU4) is responsible for decisions made on past experiences ^51^. Third, among the two enriched glutamatergic hippocampal cell types (TEGLU21 and TEGLU23), the latter expresses lipoprotein lipase as one of its top marker genes, an enzyme that causes weight gain when pharmacological or genetically attenuated in mice ^52^. Similarly, fasting inhibits activation of hippocampal CA3 cells (based on c-fos levels) in mice ^53^ and activation of glutamatergic hippocampal pyramidal neurons increases future food intake in rats most likely by perturbing memory consolidation related to the previous meal ^54^. In sum, our results provide further evidence that processes related to learning and memory play a key role in human obesity, and provide insights into specific cell types underlying hippocampal-centric susceptibility to obesity.

### Limitations of our approach

Our results should be interpreted in the light of the underlying data used to prioritize the cell types. First, the scRNA-seq data analyzed in the paper were derived from late postnatal (Mouse Nervous System dataset) and adult mice and hence did not cover developmental states and cell types of potential importance to obesity ^6^. Second, the datasets used in this work should not be regarded as complete atlases because they are likely to miss relevant cell types (from for example the paraventricular hypothalamus, nodose ganglia and adipocytes) and cell states. Third, one should keep in mind the overall assumption behind our approach, namely that in order for a disease to manifest in a given tissue or cell type the set of disease-causing genes must be active and expressed in the given tissue or cell type. In other words, our model presupposes that high/increased expression (and not decreased/lack-of expression) of a gene results in disease. Thus, our approach is not designed to detect cell types in which reduced expression of a specific gene predisposes to obesity. Fourth, our approach assumes that cell type specific patterns of gene expression are highly homologous between humans and mice. Finally, additional work is needed to further characterize the prioritized cell types during embryonal and postnatal development and relevant murine physiological regimes. While the high polygenicity of obesity and the inaccessibility of the human brain complicate approaches to further establish the enriched cell types’ relevance in human obesity, we believe that combinations of functional imaging techniques, postmortem single-nucleus analyses, enhancers to gene maps and fine-mapping of BMI GWAS loci will be crucial to better understand their role in human obesity.

### Relevance to human obesity

Despite these limitations, several lines of evidence suggest that the cell types identified herein to be enriched for BMI GWAS signal are relevant to human obesity. First, weight gain is the most pronounced side effect of subthalamic nucleus deep brain stimulation used to treat Parkinson patients ^55^, an adverse side effect that may involve the DEINH3 cell type thought to reside in the subthalamic nucleus. Second, lorcaserin (Belviq), one of the few FDA-approved anti-obesity drugs, acts on the 5-HTR2C receptor to enhance serotonin signaling. Third, at the genetic level BMI is significantly correlated with attention deficit/hyperactivity disorder (ADHD) ^56^, and growing evidence points to links between ADHD and eating disorders. For example, lisdexamfetamine (Vyvanse), a medication used to treat ADHD, is also used to treat binge eating ^57^, while the ADHD medication methylphenidate (Ritalin) is known to reduce appetite ^58^. These pharmacological observations suggest that the shared heritability of BMI and ADHD may involve pleiotropic gene variants acting through dorsal midbrain pathways. Fourth, genetic predisposition to obesity is protective to feelings of worry ^59^, supporting our findings that these two traits are potentially acting through overlapping cell types in the dorsal midbrain. Finally, BMI variants associated with BMI in a GWAS conducted in Japanese individuals enriched most highly for enhancers active in the hippocampus ^60^ and maternal obesity is associated with reduced total hippocampal volume in reduced CA3 volume in children ^61^. Together these observations support a model in which integration of sensory signals, the dopamine system and memory are likely to play key roles in humans’ susceptibility to obesity.

In conclusion, our results implicate specific brain nuclei regulating integration of sensory stimuli, learning and memory in human obesity and provide testable hypotheses for mechanistic follow-up studies. Our methodological framework provides a salient example of how human genetics data can be integrated with murine scRNA-data to identify and map components of brain circuits underlying obesity. We provide easy to use computational toolkits, CELLECT and CELLEX, which we envision will greatly facilitate future functional interpretation of common and rare variant genetic association data.

## METHODS

### GWAS

For our primary analysis we obtained BMI GWAS summary statistics performed in UK Biobank participants (N_max_=457,824) ^26^. To examine the robustness of our results to changes in GWAS cohort size, we performed secondary analyses on BMI GWAS summary statistics from two meta-analysis described in Yengo et al. (2018) (N_max_=795,640, UK Biobank and GIANT cohorts) and Locke et al. (2015) (N_max_=322,154, European subset). We note that these two studies include individuals genotyped on custom array chips (Illumina Metabochip) which violate certain assumptions of S-LDSC, but show that this has a negligible effect on results. **Supplementary Table 1** provides the full list of GWAS summary stats analyzed in this study. We used the script “munge_sumstats.py” (LDSC v1.0.0, URLs) to prepare all GWAS summary statistics. All prepared statistics were restricted to HapMap3 single nucleotide polymorphisms (SNPs), excluding SNPs in the major histocompatibility complex region (chr6:25Mb-34Mb; the term ‘SNPs’ and ‘variants’ will be interchangeably used throughout the Supplementary Notes).

### Single-cell RNA-seq datasets

For the Tabula Muris dataset (Tabula and Consortium (2018); SmartSeq2 protocol) cell types were defined as unique combinations of cell ontology and organ annotation (for example, “Lung-Endothelial_cell”) resulting in n=115 cell type annotations (of which one was defined as neuronal). For the Mouse Nervous System dataset (Zeisel et al. (2018); 10x Genomics protocol) we used the “ClusterName” option as cell type annotations (n=265, of which 214 were defined as neuronal). For the Hypothalamus dataset we constructed a cell-type compendium comprising datasets from six studies, including four covering distinct regions and two pan-hypothalamic datasets:

- ARC-ME: Arcuate nucleus and median eminence complex (from Campbell et al. (2017), DropSeq protocol). We used the “Subcluster” annotations (n=65, of which 34 were defined as neuronal).
- POA: Preoptic area (from ^33^, 10X Genomics protocol) dataset. We used the “Non-neuronal.cluster.(determined.from.clustering.of.all.cells)” annotations for non-neuronal cell types (n=21) and the “Neuronal.cluster.(determined.from.clustering.of.inhibitory.or.excitatory.neurons)” annotation for neuronal cell types (n=66).
- LHA: Lateral Hypothalamic Area (from ^32^, 10x Genomics protocol). We used the “dbCluster” annotations (n=43, of which 30 were defined as neuronal).
- VMH: Ventromedial Hypothalamus (from ^31^, SMART-seq and 10x Genomics protocols). We used the “smart_seq_cluster_label” annotations for the SMART-seq dataset (n=48, of which 40 were defined as neuronal) and the “tv_cluster_label” annotation for the 10x Genomics dataset (n=29, all defined as neuronal).
- HYPC: Pan hypothalamus (from ^34^, DropSeq protocol). We used the “SVM_clusterID” annotations (n=45, of which 34 were defined as neuronal).
- HYPR: Pan hypothalamus [from ^35^, Fluidigm C1 protocol) dataset, we used the “level1 class” annotation for non-neuronal populations (n=6) and the “level2 class (neurons only)” annotation for neurons (n=54).

Code to download and reproduce preprocessing of all datasets are available via GitHub (URLs). **Supplementary Tables 2,3,12** lists cell type annotations, the number of cells per cell type and relevant meta-data for each dataset analyzed (for the Hypothalamus dataset we list the cell-type labels used in this study as well as and the cell-type labels used in the original studies).

### Single-cell RNA-seq data pre-processing

For each dataset, we began with a matrix of gene expression values. We normalized expression values to a common transcript count (with *n*=10,000 transcripts as a scaling factor) and applied log-transformation (*log*(*x* + 1)). Next we excluded ‘sporadically’ expressed genes following the approach described in Skene *et al.* ^18^ using a one-way ANOVA with cell type annotations as the grouping factor and excluding all genes with *P*>10^-5^. We mapped mouse genes to orthologous human genes using Ensembl (v. 91), keeping only 1-1 mapping orthologs. See **Supplementary Notes** for a detailed description and discussion of the snRNA-seq data pre-processing.

### CELLEX expression specificity

See **Supplementary Notes** for a detailed discussion on ES calculations, assumptions and limitations. CELLEX version 1.0.0 was used to produce all results reported in this manuscript. A ready-to-use Python implementation of CELLEX can be retrieved from https://github.com/perslab/CELLEX. We calculated expression specificity separately for Tabula Muris, Mouse Nervous System and each of the hypothalamus datasets. Cell type expression specificity weights (*ES_w_*) were calculated using four expression specificity metrics (ES metrics) referred to us as *Gene Enrichment Score (GES)* ^24^, *Expression Proportion (EP)* ^18^, *Normalized Specificity Index (NSI)* ^62^ and *Differential Expression T-statistic (DET)*. The mathematical formulas for the ES metrics can be found in **Supplementary Notes**. For each ES metric we separately computed gene-specific *ES_w_*s before averaging them into a single ES estimate (*ES_μ_*) using the following steps:

1. For each cell type we determined the set of specifically expressed genes, *G_s_*, by testing the null hypothesis that a gene is no more specific to a given cell type than to cells selected at random. We computed empirical *P*-values of ES weights by comparing observed weights to ‘null’ weights obtained by permuting the dataset’s cell type annotations.
2. For each cell type we calculated *ES_w*_* representing the genes’ score of being specifically expressed in a given cell type. We assumed that each cell type has a set of specifically expressed genes exhibiting a linearly increasing score reflecting its expression specificity. We modeled this linearity assumption by rank normalizing *ES_w_* for genes, *g*, in *G_s_*:

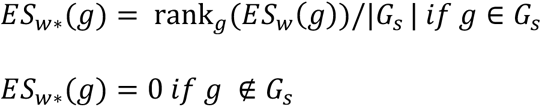

Note that *ES_w*_* are scaled such that *ES_w*_* ∈ [0,1].

3. For each cell type, we calculated *ES_μ_*, representing a gene’s score of being specifically expressed in a given cell type, by taking the mean *ES_w*_* across all ES metrics (we here assume equal weighing of ES metrics).

We use ‘*ES genes*’ to denote the set of genes with *ES_μ_*>0 for a given cell type. Hence, all genes being part of at least one *G_s_* for a specific cell-type will be included in the set of *ES genes* for this cell-type. **Supplementary Fig. 1** and **Supplementary Tables 2,3,12** show the number of *ES genes* for each cell type and dataset analyzed. We note that *ES genes* captures genes that were not only strictly specifically expressed (only expressed in the cell type) but also those that were loosely specifically expressed (i.e., have higher expression in the cell type). All cell type prioritization results were based on the *ES_μ_* estimates.

### CELLECT genetic prioritization of trait-relevant cell types

See **Supplementary Notes** for a detailed discussion on assumptions and limitations. CELLECT version 1.0.0 was used to produce all results reported in this manuscript. A ready-to-use Python implementation of CELLECT can be retrieved from https://github.com/perslab/CELLECT. Throughout this paper we report CELLECT cell type prioritization results using S-LDSC, as this model has been shown to produce robust results with properly controlled type I error ^63^. Cell type prioritization results using MAGMA ^30^ can be found in **Supplementary Fig. 5** and **Supplementary Table 10**.

#### S-LDSC

We used stratified LD Score Regression (S-LDSC) (v. 1.0.0, URLs) to prioritize cell types after transforming cell type *ES_μ_* vectors into S-LDSC annotations. Running S-LDSC with custom annotations follows three steps: generation of annotation files, computation of annotation LD scores and fitting of annotation model coefficients. We created annotations for each cell type by assigning genes’ *ES_μ_* values to genetic variants utilizing a 100 kilobase (kb) window of the genes’ transcribed regions. Fulco *et al.* showed that most enhancers are located within 100 kb of their target promoters ^64^. When a variant overlapped with multiple genes within the 100-kb window, we assigned the maximum *ES_μ_* value. The relatively large window size was chosen to capture effects of nearby regulatory variants, as the majority of trait-associated variants have been shown to be located in non-coding regions ^65^. Our results were robust to changes in window size (data not shown), consistent with previous work ^18, 63, 66^. Following the recommendation in ^63^ we constructed an “all genes” annotation for each expression dataset, by assigning the value 1 to variants within 100-kb windows of all genes in the dataset. We used hg19 (Ensembl v. 91) as reference genome for genetic variant and gene chromosomal positions. When constructing annotations, we used same 1000 Genomes Project SNPs ^67^ as in the default baseline model used in S-LDSC. Next we computed LD Scores for HapMap3 SNPs ^68^ for each annotation using the recommended settings.

For the primary cell type prioritization analysis, we jointly fit the following annotations: (i) the cell type annotation; (ii) all genes annotation (iii) the baseline model (v1.1). For cell type conditional analysis (**Supplementary Fig. 6**) we added (iv) the cell type annotation conditioned on when fitting the model.

We ran S-LDSC with default settings and the workflow recommended by the authors. We reported *P*-values for the one-tailed test of positive association between for trait heritability and cell type annotation *ES_μ_*. We note that the correlation structure among *ES_μ_* for cell type annotations can lead to a distribution of *P*-values that is highly non-uniform ^63^. Highly significant *P*-values occur due to correlated cell types with true signal, whereas cell types negatively correlated with the true signal have *P*-values near 1. For all results, we used Bonferroni correction within a trait and dataset to control the FWER. We report the regression effect size estimate for each cell type (**Supplementary Tables 6,7,12**; “Coefficient” column), which represents the change in per-SNP heritability due to the given cell type annotation, beyond what is explained by the set of all genes and baseline model. We also report standard errors of effect sizes (“Coefficient std error” column), computed using a block jackknife ^13^. Finally, we report the “annotation size” for each cell type, that measures the proportion of SNPs covered by the cell type annotation (0 means no SNPs were covered by the annotation; 1 means all SNPs were covered). Annotation size was computed as the mean of the cell type annotation.

#### S-LDSC heritability analysis

All S-LDSC heritability analysis and reported effect size estimates were obtained on the observed heritability scale, with the exception of heritability estimates for case-control traits shown in the barplots of **Fig. 2a** and **Fig. 3b**. Here we report heritability estimates on the liability scale using population prevalences listed in **Supplementary Table 1**. (The liability scale is needed when the aim of heritability analysis is to compare heritability estimates across traits. On a liability scale the case-control trait is treated as if it has an underlying continuous liability, and then the heritability of that continuous liability is quantified.) To interpret the heritability explained by our continuous-valued *ES_μ_* cell type annotations, we estimated the heritability of each *ES_μ_* quintile. We modified the script ‘quantile_M.pl’ (from the LDSC package) to compute heritability enrichment for five equally spaced intervals of the cell types *ES_μ_* annotations: (0-0.2], (0.2-0.4], (0.4-0.6], (0.6-0.8], (0.8-1], as well as the interval including zero values only ([0-0]).

#### MAGMA cell type prioritization

To assess the robustness of the SNP-level S-LDSC cell type prioritization, we used an alternative gene-level approach inspired by ^18^ and tested the association of gene-level BMI association statistics with cell type *ES_μ_* using MAGMA (v1.07a) ^30^. MAGMA was run with default settings to obtain gene-level association statistics calculated by combining SNP association *P*-values within genes and their flanking 100-kb windows into gene-level Z-statistics, while accounting for LD (computed using the 1000 Genomes Project phase 3 European panel; ^67^). Gene-level Z-statistic were corrected for the default MAGMA covariates: gene size, gene density (a measure of within-gene LD) and inverse mean minor allele count, as well the log value of these variables. Next, we used the R statistical language to fit a linear regression model using MAGMA gene-level Z-statistics as the dependent variable and cell type *ES_μ_* as the independent variable. We report cell type prioritization *P*-values (from the linear regression model) as the positive contribution of cell type *ES_μ_* regression coefficient to BMI gene-level Z-statistics (one-sided test).

### Cell type geneset enrichment

To assess cell type enrichment of genesets associated with obesity, we tested if members of the obesity geneset exhibited higher expression specificity (*ES_μ_*) in the given cell type than non-members of the geneset (all other genes in the dataset). Specifically, we used Mann-Whitney U test to obtain one-sided geneset enrichment *P*-values. We controlled the FWER using the Bonferroni method calculated over all cell types and the rare variant obesity geneset tested. As a precaution against unknown confounders, we also computed empirical p-values by permuting the expression specificity gene labels 10,000 times to obtain ‘null genesets’ of identical size, and obtained near-identical results (data not shown). We obtained genes with rare coding variants associated with obesity (n=13 genes) and genes implicated in syndromic obesity (n=39) from Turcot et al. (2018) Table 1 and Supplementary Table 21, respectively. We combined these genes into a single geneset (‘*rare variant obesity geneset*’) consisting of unique 50 unique genes.

### Cell type gene co-expression networks

We identified cell type gene co-expression networks using Weighted Gene Correlation Network Analysis (WGCNA) framework proposed by Langfelder and Horvath (2008). To identify gene co-expression networks (henceforth referred to as *gene modules*) operating within a cell type, the input to WGCNA is expression data for individual cell types. Briefly our framework consisted of the following steps:

1. We normalized the raw expression values to a common transcript count (with n=10,000 transcripts as a scaling factor), log-transformed the normalized counts (log(x+1)), and centered and scaled each gene’s expression to Z-scores. Cell clusters with fewer than 50 cells were omitted, and genes expressed in fewer than 20 cells were removed. We then used PCA to select the top 5000 highly loading genes on the first 120 principal components. We mapped mouse genes to orthologous human genes using Ensembl (v. 91), keeping only 1-1 mapping orthologs.
2. We then used hierarchical clustering and hybrid tree cutting algorithms to identify gene modules. Module eigengenes, which summarise module expression in a single vector, were computed and used to identify and merge highly correlated modules.
3. Finally, we computed gene-module correlations (kMEs), a measure of gene-module membership, filtering out any genes which were not significantly associated with their allocated module after correcting for multiple testing using the Benjamini-Hochberg method.

### Genetic prioritization of cell type co-expression networks

Genetic prioritization of WGCNA gene modules followed the same framework as for prioritizing cell types. That is, we used S-LDSC controlling for the baseline and “all genes” annotations. Gene modules annotations were constructed by assigning the module genes’ kME values to variants within a 100-kb window of the genes’ transcribed regions. We restricted modules to contain at least 10 genes and at most 500 genes (removing 8 out of 571 modules), because S-LDSC is not well-equipped for prioritizing annotations that span very small proportion of the genome, and unspecific modules with a large number of weakly connected genes have limited biological relevance.

### Co-expression networks visualizations

To create the network visualization of the cell type rWGCNA gene modules (**Supplementary Fig. 4b**), we computed the Pearson’s correlation between module kME values (a measure of gene-module membership) and generate a weighted graph between modules using the positive correlation coefficients only. To create the network visualization of the M1 gene module (**Supplementary Fig. 4c**), we computed the Pearson’s correlation between genes within the module, using expression data from the cell type in which the module was identified (MEINH2). We then generate a weighted graph between genes using the positive correlation coefficients only. We then mapped MAGMA BMI gene-level Z-statistics (calculated using 100-kb windows, as described above) onto the network as node sizes. All networks were visualized using the R package ‘ggraph’ with weighted Fruchterman-Reingold force-directed layout.

### Cell type enrichment of co-expressed gene networks

To assess if gene modules were enriched in the expression specific genes of specific cell types, we tested if module gene members exhibited higher expression specificity (*ES_μ_*) in the given cell type than non-members of the module (all other genes in the dataset). We obtained one-sided enrichment P-values using the Mann-Whitney U test. We controlled the FDR by using the Bonferroni method calculated over gene modules tested.

## ACKNOWLEDGEMENTS

Novo Nordisk Foundation Center for Basic Metabolic Research is an independent Research Center, based at the University of Copenhagen, Denmark and partially funded by an unconditional donation from the Novo Nordisk Foundation (www.cbmr.ku.dk) (Grant number NNF18CC0034900). THP acknowledges the Novo Nordisk Foundation (Grant number NNF16OC0021496) and the Lundbeck Foundation (Grant number R190-2014-3904). PNT acknowledges the Danish Ministry of Higher Education and Science for the Elite Research PhD scholarship.

We gratefully acknowledge Diego Calderon for helpful discussions on genetic prioritization models; Steven Gazal for support with LDSC heritability enrichment; Christiaan de Leeuw for support on MAGMA; Stephen Quake, Spyros Darmanis and the Biohub team for providing pre-publication access to the Tabula Muris dataset; Michael W. Schwartz, Thorkild I.A. Sørensen, Lars Ängquist and Dylan Rausch for helpful inputs on neuroendocrinology and obesity; Tobias Stanius, Ben Nielsen and Tobi Algebe for helpful discussions on expression specificity.

## AUTHOR CONTRIBUTIONS

PNT and THP designed the study and wrote the manuscript; PNT and THP conceptualized CELLECT and CELLEX, which then were developed by PNT; PNT implemented and performed all computational analyses and visualizations except for WGCNA analysis performed by JJT; PNT and THP analyzed and interpreted the resulting data. All authors read and approved the manuscript.

## CODE AVAILABILITY

### Software

CELLECT toolkit is available at https://github.com/perslab/CELLECT. CELLEX is available at https://github.com/perslab/CELLEX. Open source software implementations of CELLECT and CELLEX will be made available upon publication.

### Code to reproduce results and Fig

Code to reproduce analyses, Fig. and tables for this manuscript is available at https://github.com/perslab/timshel-bmicelltypes

## URLS

- LDSC: https://github.com/bulik/ldsc
- MAGMA: https://ctg.cncr.nl/software/magma
- Robust WGCNA pipeline: https://github.com/perslab/wgcna-toolbox

