## Supplementary Information for "Mapping heritability of obesity by brain cell types"

### Supplementary Note 1
### CELLECT cell type prioritization

#### Introduction

In this Supplementary Note we discuss the detailed methods and limitations of the CELLECT framework.

#### The relevance of using mouse scRNA-seq datasets

Our BMI cell type prioritization analysis was performed using mouse scRNA-seq data, as there are currently no comprehensive human multi-organ and multi-brain area cell type atlases available. We here discuss the relevance of using mouse scRNA-seq datasets to define cell types for genetic prioritization for complex human traits.

Previous studies have compared the conservation of tissue gene expression. Brawand *et al.* [1] used RNA-seq expression data across multiple organs (cortex, cerebellum, heart, kidney, liver, and testis) and 10 mammalian species (incl. human) and found the largest amount of variation was explained by organ rather than species differences.

Previous studies have assessed the convergence of mouse and human CNS gene expression using gene co-expression analysis [2–4] and found weaker conservation of glial co-expression modules than neuronal co-expression modules. *In situ* hybridization studies have reported that the majority of genes (79%) showed similar cortical laminar patterning [5]. Along those lines, recent scRNA-seq data from mouse and human midbrain found that cell types and gene expression levels were generally conserved across species [6].

The current most extensive study of CNS cell type conservation compared single-nuclei expression data from human and mouse cerebral cortex [7] and found that broadly defined cell types were conserved between mouse and human. However, they identified important differences between cell type proportions and expression of specific genes, including cell type marker genes exhibiting up to 10-fold expression differences.

In conclusion, although critical differences between mouse and human CNS gene expression data have been identified, the broad expression patterns are likely to be conserved. Moreover, glial cell types are more likely to exhibit weaker conservation compared to neuronal cell types. We believe our genetic prioritization of likely etiologic cell types is more likely to suffer from false negatives (cell types not prioritized because of lack of relevant human expression data) rather than false positives (spuriously enriched cell types among our positive results).

#### Choice of window size and position for connecting SNPs and genes

An important step in the CELLECT pipeline is assigning gene $ES_\mu$ values to SNPs. As the majority of trait-associated SNPs are located in non-coding regions [8], it is desirable to select a window size that maps the majority of regulatory GWAS variants to their proximal genes. Although our results were largely robust to changes in window size and consistent with previous work [9–11], we note that the

SNP-to-gene mapping remains a critically important step that should be updated in subsequent versions of CELLECT.

In this work, we used the same window size as used in Finucane et al. (2018), that is, assigning genes' $ES_{\mu}$ values to SNPs within a 100 kb window of the genes' transcribed regions. A recent large eQTL analysis in blood from >31,000 individuals found that 92% of the lead cis-expression quantitative trait loci (eQTL) SNPs mapped within 100 kb of the gene [12], suggesting that our mapping is likely to capture the majority of cis-regulatory variants. Consistent with this, Gasperini et al. (2018) used CRISPR/Cas9 followed by scRNA-seq to identify CRISPR/Cas9-induced eQTLs from >47,000 human cell line cells and found that regulatory variants were separated from the TSS of their target genes by a median distance of 34.3 kb. Finally, work by Fulco *et al.* reports that most enhancers are located within 100 kb of the target promoters [14].

### Limitations of CELLECT

#### Linear relationship between expression specificity and trait heritability

The overall assumption behind our approach is that in order for a disease to manifest in a given cell type the set of disease causal genes must be active and expressed in the given cell type. In other words, we assume that high/increased expression and not decreased/lack-of expression of a gene results in disease. This is a strong assumption to make about complex traits and it does not hold for all diseases (e.g. cancer).

Our model assumes a linear effect of cell type expression specificity and trait heritability. Although this assumption may not always hold, it appears to be reasonable in the continuous annotations that we analyzed (**Supplementary Fig. 11a**).

We leave it for future work to explore non-linear relationships between expression specificity and trait heritability, and to investigate the effects of specificity for decreased or lack-of gene expression.

#### Genetic architecture

The approach assumes that a cell type is etiologic for a particular disease if and only if genetic variants near genes with high expression specificity in the cell type are enriched for heritability. Moreover, the CELLECT cell type prioritization assumes a polygenic trait architecture. Consequently, our approach is unlikely to yield relevant results for traits driven by rare genetic mutations (not covered by GWAS) or traits where the heritability is not mediated by transcriptional differences (i.e. changes related to other molecular modalities such as proteins, posttranslational modifications or the microbiome).

#### Common variation

We restricted our analysis to common variants (HapMap3 SNPs, >5% MAF), as S-LDSC has several limitations when applied to rare variants. Prioritizing cell types using a model that includes both common and rare variants could produce different results. We argue that our results are likely to be robust to changes in the allele frequency spectrum. Firstly, we found that cell types enriched for rare variant obesity genes overlapped with the S-LDSC BMI prioritized cell types (based on common variants). Secondly, a recent study by Zhu and Stephens (2018) compared the ability of genetic enrichment methods (incl. S-LDSC) to detect the true enrichment signal based on 1000 Genomes Project SNPs and HapMap3 SNPs, and found that all methods (S-LDSC included) produced similar results using the two sets of SNPs as input. Thirdly, rare variants are unlikely to explain the majority of BMI heritability: Gazal *et al.* estimated the low-frequency variants (MAF<5%) to explain 15% of BMI heritability [16] and recent work (*under review*) has reported that variants with MAF<10% might explain as much as 51% of BMI heritability [17]). In conclusion, cell type prioritization results restricted to common variants are likely to converge with results including rare variants.

#### Expression heritability mediated by cis- vs trans-eQTLs

As our model assumes that SNPs *near* genes with high expression specificity in etiologic cell types are enriched for heritability, our model relies on the majority of gene regulatory variants (eQTLs) are located nearby (*cis-acting*) instead of distant (*trans-acting*) to the target gene. That is, we assume that heritability of gene expression can be sufficiently explained by *cis*-acting variation. There are notable examples where the causal regulatory variant act *in trans*, e.g. the causal variant located in the first intron of the FTO locus are located >1 Mb from its target regulatory genes IRX3/IRX5 [18,19], but the question how prevailing *trans*-acting variation is remains unresolved. In support of the sufficiency of *cis*-variation, one study found that *cis*-eQTLs explain a substantial proportion of trait heritability (40-80%) [20]. In addition, transcriptome-wide association studies (TWAS) leverage *cis-eQTLs* to predict expression levels with a 60-80% prediction accuracy [21,22]. In contrast, Liu et al. (2019) report that up to 60-90% of genetic variance in expression is due to trans-acting variation. We acknowledge that trans-acting effects are likely to play an important role in gene expression heritability, but despite promising efforts [14] cell type-specific enhancer to gene maps have not been constructed yet and hence we based CELLECT on cis-regulatory variants only.

### Future directions

We envision several improvements of our approach. SNP-to-gene mapping could be improved by levering for instance the ABC model propsed by Fulco *et al.* [14] cell types to predict enhancer to promoter maps to assign regulatory variants to genes. Alternatively, SNP-to-gene mapping could be improved by using LD-informed loci definitions centered on SNPs. That is, each SNP would be assigned $ES_{\mu}$ value based on the genes within the LD defined loci boundaries of the SNP (e.g. genes within the region spanned by $r^2 < 0.7$).

It would be of interest to explore non-linear relationships between expression specificity and heritability and to test whether down-regulated genes contribute to cell type heritability. Such analysis would be possible leverage an extension of LDSC referred to as signed linkage disequilibrium profile regression [24], which allows detection of directional effects of signed functional annotations.

Finally, we envision a data-driven approach to select the parameters of our approach, e.g. SNP-to-gene parameters or non-linear transformation of expression specificity. A genetic trait with known etiologic cell types could be used to select the set of parameters resulting in the most significant prioritization of the known etiologic cell types.

### Supplementary Note 2
### Omnigenic model notes

#### Introduction

In this Supplementary Note we attempted to unify our cell type prioritization model with the so-called omnigenic hypothesis proposed by Prichard and colleagues [23,25].

#### Levering the omnigenic model to detect disease causal cell types

In the single-cell era, we are for the first time able to leverage the power of the proposed omnigenic genetic architecture, by identifying cell type specific expression of core and peripheral genes. We here describe the key assumptions and approach behind CELLECT.

##### Key assumptions and observations

Our key assumption is that in order for a disease to manifest in a given tissue or cell type the set of disease causal genes must be active and expressed in the given tissue or cell type. That is, we assume that high/increased expression and not decreased/lack-of expression of a gene results in the given trait or disease (henceforth simply referred to as *disease*). This is a strong assumption to make about complex traits and it does not hold for all diseases (e.g. cancer).

We assume that for a cell type to be causal to a given disease, it should express **one or more core genes**. We note an important distinction between this assumption and the stronger more commonly used assumption that causal, disease cell types enrich for expression of all core genes (e.g. testing for top expressed cell type genes for enrichment of genes harboring rare variants [10]. Because core genes can function in orthologous pathways, we only assume expression of one or more core genes.

We assume that **core genes have cell type specific etiologic roles** for common complex traits. This assumption is justified by the strong negative selection of mutations in genes with a ubiquitous function broadly affecting cellular function. (Detrimental mutations in genes with non-redundant basic cellular function will not manifest in the population as a common disease.) We note that this only holds true for *heritable common* diseases. E.g. core driver cancer genes may have basic biological functions as observed with *de novo* mutations in *TP53*.

We reason that the majority of genes localizing near GWAS loci explaining the most heritability are **peripheral genes** that **are more likely than other genes to exhibit cell type-specific expression**. This assumption is justified by two steps of reasoning. Firstly, **peripheral genes can only exert their effect on core genes if they are co-expressed in a cell.** They must operate within the same network of expressed genes in a cell (see ref. [25] Fig. 4b). Secondly, GWAS is more well-powered to detect peripheral genes in close proximity to core genes, if a shorter degree of separation between peripheral and core genes increases the effect of the peripheral gene on the core gene (*ibid.*; Fig. 4a). We note that under the 'small world' network property of gene regulatory networks, most expressed genes in a cell type are only a few steps from the nearest core gene, possibly making the set of 'peripheral genes in close proximity to core genes' quite large.

**Biological examples of potential mechanisms**

It has been shown that impaired signaling from the primary cilia of *MC4R-positive* neurons can cause obesity in humans. *In vitro* and *in vivo* work from Siljee et al. (2018) demonstrated that *MC4R* obesity-causing mutations impair the localization of *MC4R* to the cilia. The discovery of obesity-causal mutations in *ADCY3* provided evidence that *ADCY3* plays a role in human obesity [27,28]. This example may demonstrate how genetic variation in one peripheral gene, *ADCY3*, can regulate/impair the function of a core gene, in this case, *MC4R*, within the energy-regulating melanocortin signaling pathway. The omnigenic model suggests that there are many yet unknown genetic regulators affecting *MC4R* (potentially also through primary cilia signaling or targeting) and hence contributing to the obesity heritability.

Our results for prioritized cell types could support this distinction between core and peripheral gene. We find that the core gene (*MC4R*) is co-expressed with the peripheral gene (*ADYC3*) (**Fig. S2.1**). *MC4R* is highly specifically expressed and *ADCY3* is moderately specifically expressed.

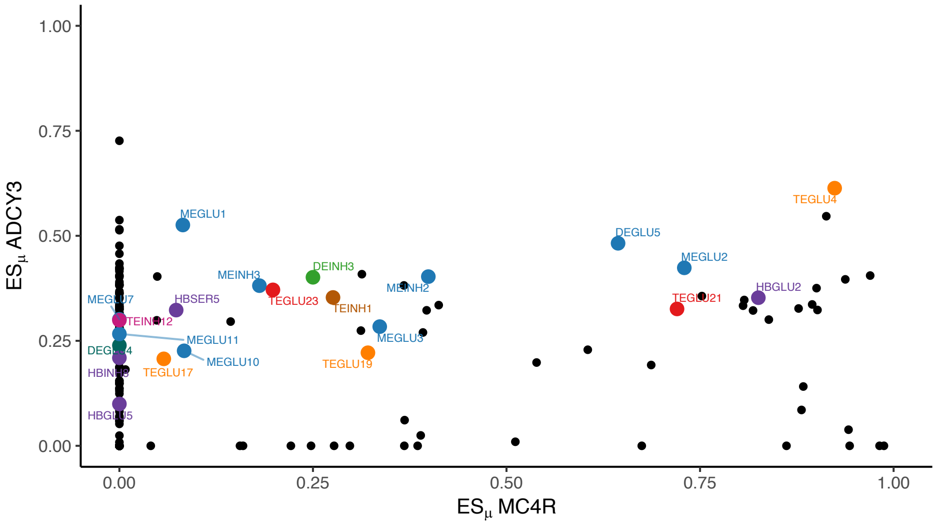

**Fig. S2.1 | Co-specific expression of *MC4R* and *ADCY3*.** BMI-prioritized cell types highlighted in color. TEGLU4 (located in cortex pyramidal layer 5) is a candidate etiologic cell type mediating the role of MC4R/ADCY3 in obesity.

### The model

Our framework, CELLECT, is to the best of our knowledge, the first method to use *continuous* LDSC annotations to identify likely disease causal cell types. We here explain why using *continuous* (i.e. weighted) annotations is an important improvement over existing studies.

**We assume that if a gene is specifically expressed in a given cell type, it is functionally important for that cell type.** That is, specifically expressed genes constitute the functionally distinct part of the cell type. For instance, we have shown that *DRD2* is specifically expressed in certain cell types from the midbrain, suggesting the functional role of these cell types in the dopamine reward system. *ES genes* will, by definition, not contain ubiquitous/equally expressed genes (e.g. basic cellular/biological processes). Please also refer to **Supplementary Note 3**.

Following our above assumptions, for a disease causal cell type we assume the following relationship of cell type specific expression:

$$ES(trait\ relevant\ core\ genes) \geq ES(trait\ peripheral\ genes) > ES(non\ trait\ relevant\ genes)$$

We are now able to express our model for genetic identification of causal disease cell types. Formally we model a linear relationship between a gene's disease heritability and cell type expression specificity:

$$Heritability(gene) \sim ES(gene)$$

**Fig. S2.2** shows the concept of how the two above equations can be used to identify likely disease causal cell types.

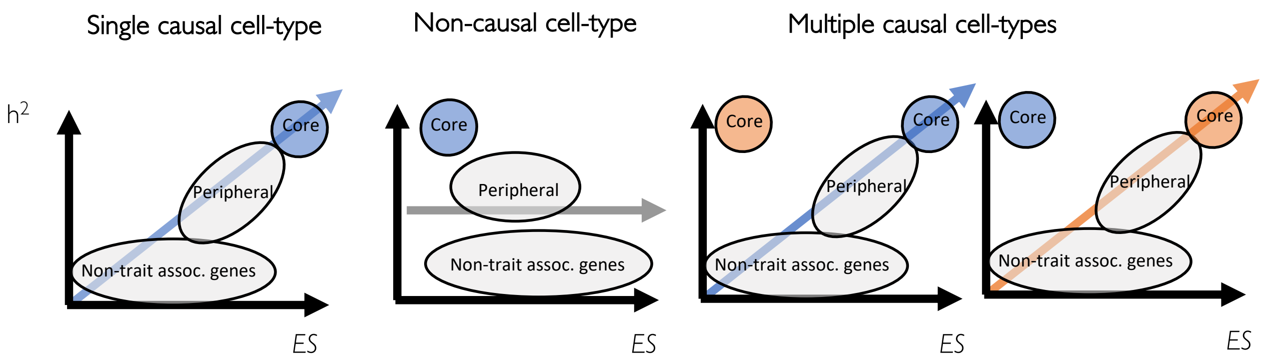

**Fig. S2.2 | Levering the omnigenic model to identify causal cell types using expression specificity**. A linear model can be used to identify causal cell types by testing for association between gene expression specificity and gene heritability. Three scenarios are shown: a causal cell type (left), a non-causal cell type (middle) and multiple causal cell types (right).

In support of this model, we showed that top expression specific genes in BMI-prioritized cell types exhibit higher BMI heritability enrichment than non-expression specific genes (**Supplementary Fig. 11a**).

### Summary

We here provide a brief summary of the points discussed in the above sections.

#### Model assumptions and observations

- The "omnigenic" genetic architecture of complex traits states that so-called peripheral genes (peripheral referring to the core molecular function encoded by the *core* genes in the cell type) explain the majority of heritability, and as a consequence, identifying heritability enrichment using only core genes, will fail.
- To identify disease causal cell types, we estimate the disease heritability explained by cell type specifically expressed genes.
- Likely disease causal cell types have one or more core genes specifically expressed *and* the specifically expressed genes enrich for disease heritability.
- Core genes have cell type specific roles and are cell type specifically expressed.
- Peripheral genes are co-expressed with at least one core gene in the disease-causal cell type.

- Peripheral genes are more likely than other genes to be cell type specifically expressed.
- Many peripheral genes are shared among related traits. This leads to a partial overlap between prioritized cell types for related traits. Core genes have little overlap between related traits.

#### Results

- We show that all our prioritized cell types express one or more 'core' obesity genes (as defined by our rare variant obesity geneset). See **Supplementary Fig. 8**.
- We show that top expression specific genes in prioritized cell types have higher BMI heritability enrichment than non-specifically expressed genes (**Supplementary Fig. 11a**).
- We find that the BMI prioritized cell types are, to some degree, trait specific (**Supplementary Table 5**). This implies that CELLECT is not only prioritizing shared peripheral genes between brain-related traits.

### Supplementary Note 3
### CELLEX expression specificity

#### Introduction

At the core of using expression data for genetic identification of cell types underlying disease, lies the problem of finding a meaningful vector representation of cell type expression profiles. In our approach, we represent cell types by their expression specificity (ES) profile: a measure of relative gene expression levels. To robustly estimate ES, we developed a computational method called CELLEX (**Cell type EX**pression-specificity).

In this Supplementary Note we provide additional details and limitations of the CELLEX expression specificity framework. We provide a comprehensive benchmark of ES metrics on single-cell RNA-sequencing (scRNA-seq) and show that our combined metric, $ES_\mu$, is most robust than single expression specificity measures. For an ES metric benchmark on bulk data we refer to Kryuchkova-Mostacci and Robinson-Rechavi (2016).

#### CELLEX expression specificity

##### Overview of ES computation

##### Notation

We use the term 'ES metric' to describe the given metric used to compute $ES_w$ (see **Table S3.1** for an overview of the ES metrics used). $ES_w$ are gene-level statistics computed for a given ES metric. $ES_{w^*}$ are the genes' likelihood of being specifically expressed given the ES metric. $ES_\mu$ are the genes' marginal likelihood of being specifically expressed. We use '*ES genes*' to denote the set of genes with $ES_\mu$>0 for a given cell type.

ES values are computed separately for each dataset. **Fig. S3.1** provide an overview of the steps.

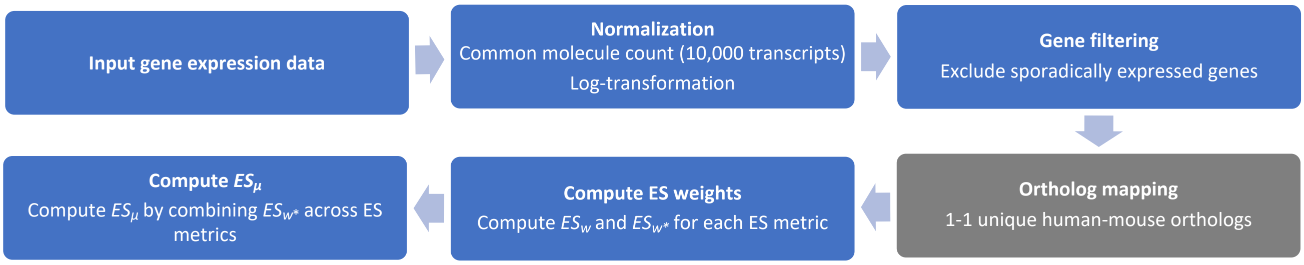

**Fig. S3.1 | Overview of CELLEX expression specificity estimation**.

##### Expression data pre-processing and normalization

We use unnormalized gene expression values as input for ES calculations. We note that the ES metrics used in this work do not allow for adjusting out unwanted sources of variation between cell type (e.g. batch effects). We assume that there are no major confounding factors between cell types.

We scale expression values to a common transcript count with n=10,000 transcripts as a scaling factor. Next, we apply log-transformation ($x_{log} = log(x + 1)$). We apply this normalization procedure for the following three reasons:

1) Common transcript count makes expression values comparable among cells.
2) Log-normalization is a variance-stabilizing transformation that dampens the effect of the dynamic range. It transforms the expression data into a more well-behaved distribution that is better approximated by a normal distribution (we later compute ANOVA and t-statistics that assume normality of the data).
3) This normalization procedure is a common, robust and proven-to-work method for scRNA-seq (it is the default normalization technique for standard single-cell analysis packages such as Seurat; Satija et al., 2015). This normalization method was also used by the authors of the two primary data sets analyzed in our study: the Mouse Nervous System [31] and Tabula Muris [32] datasets.

Despite the above strengths, we note two limitations of this normalization approach:

- When estimating the common transcript count "scaling factors" for each cell, we assume each cell has the same number of total molecules. This is an overly simplistic assumption as cell size is generally correlated with the amount of mRNA in the cell.
- Log-transformation is an empirical choice for variance-stabilizing transformation. Recently developed generalized linear models might provide better normalization [33,34].

We leave it for future work to explore additional normalization procedures and approaches to correct for confounding factors between cell types prior to estimating ES weights.

#### Gene filtering

We observed that the ES metrics were prone to falsely estimating genes with 'sporadic' gene expression levels as highly expression specific. We excluded these genes to reduce bias in the null expectation for $ES_w$ and hence reduce false positive ES genes. Most notably, EP would estimate genes with very low expression appearing in few number of cells as highly expression specific. To solve this problem, we sought to estimate the background noise level for each gene, enabling us to distinguish between genes with undetectable sporadic expression levels and genes with confident expression levels. Following the approach described in Skene et al. (2018), we reasoned that genes with sporadic expression would fail to be statistically differentially expressed in at least one cell type. We modelled this using one-way ANOVA with cell type annotations as the grouping factor and excluded all genes with $P > 10^{-5}$.

#### Invariance to gene filtering

The gene filtering and mouse-human ortholog mapping steps considerably reduces the number of genes in the dataset. We computed ES weights *after* these gene filtering steps. All ES metrics except EP (Expression Proportion) were invariant to the 'gene universe' of the dataset (all genes in the final dataset), consequently $ES_w$ are generally robust to gene filtering operations. However, $ES_{w^*}$ (denoting a genes likelihood of being specifically expressed) is sensitive to the 'gene universe' as we use a null distribution pooled across genes to determine $ES_w$ significance (see the below section "Determining the set of ES genes for each ES metric" for details).

#### Choice of ES metrics

To implement a 'wisdom of the crowd' approach, we aimed at combining a diverse set of ES metrics. We choose ES metrics based on their documented evidence to identify tissue expression specificity based on bulk expression data [29] or if they had been successfully applied on scRNA-seq datasets. Lastly, we aimed for metrics to estimate over-expression and not under-expression compared to other cell types.

Although we incorporated a t-test for differential expression as part of our ES metrics, we reasoned that test statistics or *P*-values from differential expression tests – e.g. DESeq2 [35], MAST [36] – were not sufficient for making a useful ES metric. As a result, we did not chose to build $ES_{\mu}$ purely from DE test statistics. The result of DE tests was a list of genes ranked by the *uncertainty* or *signal-to-noise ratio* (*P*-value) of the estimate for difference in expression between cell type populations. For instance, a gene exhibiting a subtle difference in expression between two cell types and very low expression variance, would result in a highly significant DE P-value. A useful ES metric should not be biased towards assigning high expression specificity to identifying genes with low variance, in part because the variance of gene expression is confounded by the cell type clustering resolution. Cell types with high heterogeneity will have higher variance of expression, resulting in downward-biased estimates of expression specificity for these cell populations when using a DE test as ES metric. In summary, our ES metrics do not seek to capture the uncertainty of difference in expression, but instead the relative magnitude of difference in expression between cell types.

#### Overview of ES metrics used in this study

We here provide a short overview of ES metrics and their interpretation. In the following table we use *c* to denote the focal cell type and *g* to denote the gene for expression specificity estimation.

| **ES metric abbr.** | **ES metric name** | **$ES_w$ scale** | **$ES_w$ interpretation** | **Reference** |
|---|---|---|---|---|
| GES | Gene Enrichment Score | $\mathbb{R}_{\geq 0}$ | <1: *g* depleted in *c*<br><br>1: *g* no enrichment<br><br>>1: *g* enriched in *c* | Zeisel (*Cell*, 2018) |
| EP | Expression Proportion | [0, 1] | 0: *g* not expressed in *c*<br><br>0.5: *c* makes up 50% of *g* total mean expression.<br><br>1: *g* uniquely expressed in *c* | Skene (*Nature Genetics*, 2018) |
| NSI | Normalized Specificity Index | [0, 1] | 0: *g* not expressed in *c*<br><br>0.5: *g*'s mean expression fold-change (focal cell type compared to other cell types) is on average within the top 50% of all genes<br><br>1: *g*'s mean expression fold-change is the largest fold-change observed over all genes. | Modified from Dougherty (*Bioinformatics*, 2010) |
| DET | Differential Expression T-statistic | $\mathbb{R}$ | <0: *g's* mean expression lower in *c*<br><br>>0: *g's* mean expression higher in *c* | - |

**Table S3.1 | ES metrics used in CELLEX.**

#### ES metrics formula

For all formulas, we calculate $\mu_{g,c}$ as the average expression of gene $g$ in cell type $c$ in a dataset with $C$ cell types and $G$ genes.

#### Gene Enrichment Score

GES is computed as the fold-change weighted by the fraction of cells expressing the gene. Zeisel et al. (2018) showed that this measure was effective at identifying marker genes for cell types in the nervous system.

$$GES_{g,c} = \frac{\mu_{g,c}}{\mu_{g,\bar{c}}} \frac{f_{g,c}}{f_{g,\bar{c}}}$$

Here $\mu_{g,\bar{c}}$ is the average expression of all other cell types, $f_{g,c}$ is fraction of cells in cell type $c$ with non-zero expression of gene $g$, and $f_{g,\bar{c}}$ is the fraction of cells with non-zero expression in all other cell types.

#### Expression Proportion

EP is calculated by dividing the expression of each gene in each cell type by the total expression of that gene in all cell types. Intuitively EP estimates the proportion of $g$'s total mean expression contributed by cell type $c$.

To remove the effect of differences in total expression between cell types, Skene et al., (2018) first normalize $\mu_{g,c}$by the total expression of cell type:

$$\mu^*{}_{g,c} = \frac{\mu_{g,c}}{\sum_{g'}^{G} \mu_{g',c}}$$

EP is then calculated as:

$$EP_{g,c} = \frac{\mu^*{}_{g,c}}{\sum_{c'}^{C} \mu^*{}_{g,c'}}$$

We note that the first normalization step has no effect on our results, since we use common molecule count normalized expression data.

#### Normalized Specificity Index

The Normalized Specificity Index (NSI) is modified from Specificity Index [37]. Intuitively NSI estimates the average quantile (or relative rank) of gene $g$'s mean expression fold-change across all cell types. NSI is given by the formula:

$$NSI_{g,c} = \sum_{k\neq c}^{K} \frac{rank_g\left(\frac{\mu_{g,c}+\epsilon}{\mu_{g,k}+\epsilon}\right) - 1}{G-1} / (k-1)$$

Where $rank_g(\mu_{g,c} \ / \ \mu_{g,k})$ gives the position of gene $g$ in a descending-ordered list of 'fold-change' values for all genes, $\epsilon$ is a small numerical constant to prevent the fold-change from going to infinity as the denominator of the fold-change goes to zero.

We modified the SI metric from ref. Dougherty et al., (2010) for two reasons. Firstly, SI was originally developed for bulk gene expression data, which makes it less well-suited for count-based scRNA-seq, which is comprises an inflation of zero values. Secondly, SI values may take on arbitrary large values depending on the number of input genes, which makes the resulting SI values hard to interpret. We sought to normalize the SI scale to an intuitive [0-1] scale. Specially, we made three relevant modifications:

1) NSI is scaled by the number of genes to obtain a [0-1] scale.
2) NSI resolve ties using the minimum value instead of the average. This is relevant for sparse scRNA-seq where many genes will be tied for zero values.
3) NSI contains a small constant, $\epsilon$, to prevent SI from going to infinity as the denominator of the fold-change goes to zero.

We used the *specificity.index()* function of the pSI R package (v1.1, release 2014-01-30) as comparison for our modifications.

#### Differential Expression T-statistic

We compute the t-statistic for gene $g$ as a measure of differential expression between the cell type $c$ and all other cell types.

$$DET_{g,c} = \frac{\mu_{g,c} - \mu_{g,\bar{c}}}{s_g\sqrt{1/n_c + 1/n_{\bar{c}}}}$$

Here, as above, $\mu_{g,\bar{c}}$ is the average expression of all other cell types, $n_c$is the number of cells in cell type $c$, $n_{\bar{c}}$ is the number of cells in all other cell types, and $s_g$ is the pooled standard deviation estimate for gene $g$ given by:

$$s_g = \sqrt{\frac{\sum_{c\prime}^{C}(n_{c\prime} - 1)s_{c\prime}{}^2}{\sum_{c\prime}^{C} n_{c\prime} - 1}}$$

#### Determining the set of ES genes for each ES metric

A key step in our approach is determining the set of *ES genes* for each ES metric. For each cell type we determined the set of specifically expressed genes, $G_s$, by testing the null hypothesis that a gene is more specific to a given cell type compared to cells selected at random. We compute empirical *P*-values of $ES_w$ by comparing observed weights to 'null' weights obtained by permuting the dataset's cell type annotations.

#### Null distribution

We use an empirical null distribution because $ES_w$ for GES, NSI and EP do not have analytic statistical distributions to assess their significance. In addition, the analytical distribution for DET is not well-calibrated for (genome-wide) single-cell DE tests.

To construct our empirical null distribution we shuffled cells corresponding to the null hypothesis: gene X is equally specific to cell type A as it is to randomly selected cells. We constructed the null distribution such that our 'null cell types' had the same number of cells as the observed cell types. We note that in our null distribution, we kept the 'cell entities' and only the genes' average expression will change. Side remark: an alternative approach to constructing the null distribution, would be to shuffle genes corresponding to the null hypothesis that gene A is not more specific for cell type X than randomly selected genes. However, this null variant is not 'scale resistant' and will hence not work if genes are not on the same normalized scale.

#### Nominal significance cut-off

We use an empirical $P$-value to find the cut-off between expression specific and non-expression specific genes. We use nominal significance ($P$<0.05) for this p-value (instead of an FDR adjusted cut-off) because we assume that our method is robust to inclusion of false positive ES genes.

#### Normalization assumptions for computing $ES_{w^*}$

For each cell type we calculate $ES_{w^*}$, representing the genes' likelihood of being specifically expressed in a given cell type and for a given ES metric, by rank normalizing $ES_w$ for genes in $G_s$:

$$ES_{w*}(gene) = rank_g(ES_w(gene))/|G_s| \; if \; gene \in G_s$$
$$ES_{w*}(gene) = 0 \; if \; gene \notin G_s$$

We set $ES_{w^*}$=0 for non-expression-specific genes (non-ES *genes*). This is important because we assume that we cannot meaningfully distinguish *between* non-ES genes, and hence they should all be given the same value.

Importantly, the rank normalization corresponds to the assumption that each cell type has a set of expression specific genes exhibiting a *linearly* increasing likelihood of being expression specific to the cell type. We apply this transformation to ensure $ES_{w^*}$ have the same scale before combining into $ES_{\mu}$.

We note that this normalization strategy discards the 'magnitude' of the expression specificity (see below section "Expression specificity profiles for ES metrics and average expression" for consequences of this). The linearity assumption essentially 'smooths' the non-linearity of $ES_w$ into a linear $ES_{w^*}$ scale. We leave it for future work if other normalization strategies (e.g. inverse normal transformation or min/max) could offer an even better trade-off between dynamic range of expression specificity and robustness of the normalization.

#### Expression specificity profiles for ES metrics and average expression

As discussed above, the magnitude of expression specificity is not captured in $ES_{w^*}$ and consequently $ES_{\mu}$. To illustrate this, we plotted expression specificity profiles for two genes from the Mouse Nervous System dataset (**Fig. S3.2**). The AGRP gene shows the same expression specificity profiles for ES metrics and log-transformed mean expression; only few cell types have high average expression levels (several order of magnitudes higher than the other cell types). The POMC gene exhibits a slightly different expression specificity profile as it is more ubiquitously expressed. For both genes, the cell types with the highest average expression were also the top expression specific genes across most ES metrics.

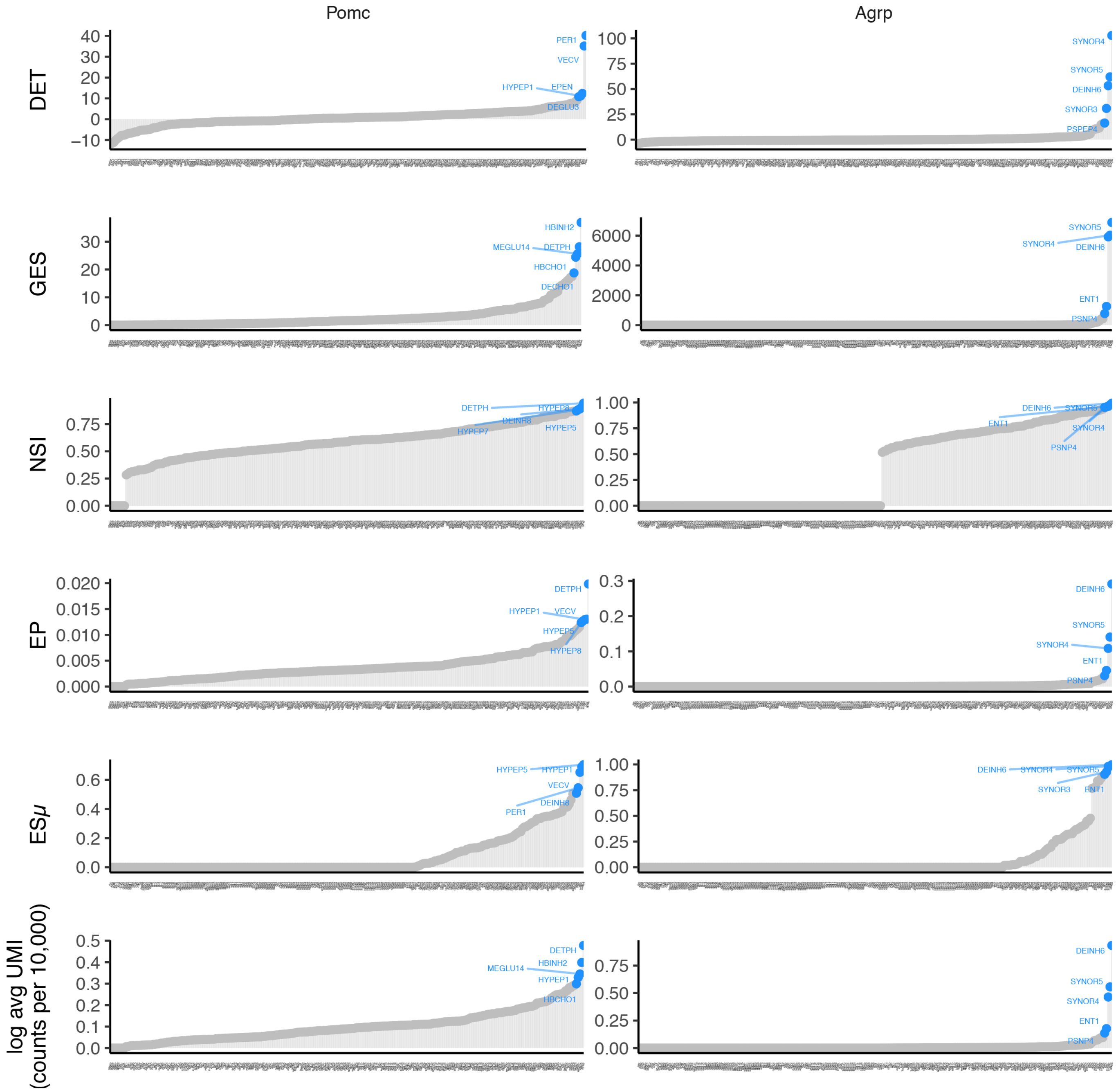

**Fig. S3.2** | Expression specificity profile for *POMC* (left) and *AGRP* (right). AGRP is expressed in few cell types and *POMC* is expressed in slightly more cell types. Each panel row shows a different ES metric (normalized average expression is shown in the bottom row panels). The plot shows gene $ES_w$ (y-axis) for each cell type (x-axis, ordered by increasing values of $ES_w$). The five cell types with the largest $ES_w$ are highlighted. $ES_w$ values are estimated from the Mouse Nervous System dataset.

#### Limitations of ES

Expression specificity is a relative measure as it depends on the background compendium of cell types included in the dataset. This means that we would obtain dramatically different ESμ values for e.g. a neuronal cell type computed on the full Mouse Nervous System dataset and on a subset consisting only of the neuronal cell types. As a consequence, ESμ values for similar cell types in two different datasets might be difficult to compare if they are computed on different background cell type compendia. Because expression specificity is inherently context dependent, there is a need for a 'common reference' dataset to ensure uniformity of values. We leave this for future work to explore.

#### Future directions

**Preprocessing of ES**: Gene expression imputation methods could be used to alleviate drop-out effects (missing values) causing downward bias of the cell type average expression estimates.

**Alternatives to rank $ES_{w^*}$ normalization**: We expect that other normalization strategies could offer a better trade-off between dynamic range of expression specificity and robustness of the normalization.

**Explore additional ES metrics**: We find it unlikely that the four ES metrics used in this work captures all aspects of cell type expression specificity. It would be highly relevant to explore additional ES metrics.

**Context dependence of ES**: We envision two strategies to combat the context dependency of ES values. One solution is to calculate a 'dataset diversity score' that measures diversity of the background compendium of cell types included in the dataset. This score could then potentially be taken into consideration when interpreting the results. A second solution could be including a 'common reference' dataset as background compendium of cell types. For example, use Seurat data integration methods [38,39] to align the Tabula Muris dataset with the dataset of interest.

### Expression Specificity robustness analysis

#### Aim

ES is inherently dependent on the context of cell types used to estimate ES values, that is the cell type composition of the dataset. Still, the $ES_{w^*}$ should primarily reflect the properties of the cell type and not the context of the dataset. For example, we wish to obtain similar $ES_{w^*}$ when we replicate a scRNA-seq experiment even if the cell type composition has shifted slightly. In other words, it is desirable for an ES metric to produce similar $ES_{w^*}$ in varying contexts of cell type composition. We here aim at assess this characteristic: the *robustness* of ES metrics. A robust metric will yield similar results in changing cell type contexts.

#### Methods

To assess the robustness of ES metrics, we defined a Robustness Score (RS), which measures the ability of an ES metric to reproduce $ES_{w^*}$ in changing cell type computations. We used $ES_{w^*}$*-baseline* to denote $ES_{w^*}$ of a selected focal cell type computed on the full dataset, *i.e.* all cell types. Further, $ES_{w^*}$*-subset* denotes $ES_{w^*}$ of the focal cell type computed on a random subset of the data. For a given focal cell type and data-subset-proportion, we performed the following procedure:

1) Create a subset of the dataset by randomly sampling the specified subset proportion (e.g. 20%) of cell types from the full dataset. The focal cell type is excluded during sampling.
2) Add the focal cell type to the sub-sampled dataset.
3) Compute $ES_{w^*}$ for the focal cell type to obtain $ES_{w^*}$*-subset*.
4) Compute Pearson's correlation coefficients between $ES_{w^*}$*--baseline* and $ES_{w^*}$*--subset* to obtain *RS-subset*.

The sampling procedure was repeated 100 times for each ES metric and data subset proportion. When computing *RS-subset*, we adjusted for zero-inflation by removing genes with zero values in both $ES_{w^*}$*-baseline* and $ES_{w^*}$*-subset*. We reported RS values averaged over the 100 repetitions.

We performed the above procedure using 12 distinct cell types as focal cell types (ABC, ACNT1, ACTE1, COP1, DEINH3, EPMB, EPSC, MGL1, OPC, PVM1, TEGLU1, VLMC1). To ensure the generalizability of our results, the 12 focal cell types were selected as representatives of each major cell class in the Mouse Nervous System dataset (astrocyte, ependymal, glia, neuronal, oligodendrocyte and vascular cells). We summarized the results of each subset proportion across the 12 focal cell types by computing the mean and standard deviation of the mean RS across cell types (**Fig. S3.3**).

#### Results

We observed that $ES_{\mu}$ achieved the highest mean RS with low variation across all tested focal cell types (**Fig. S3.3**). GES exhibited the lowest mean RS with high variation across focal cell types. The mean RS across all ES metrics was generally high (>0.8) for subsets greater than 10% of all cell types. All ES metrics exhibited similar mean RS for subsets greater than 50%. In summary, $ES_{\mu}$ is the most robust ES metric to changes in cell type composition.

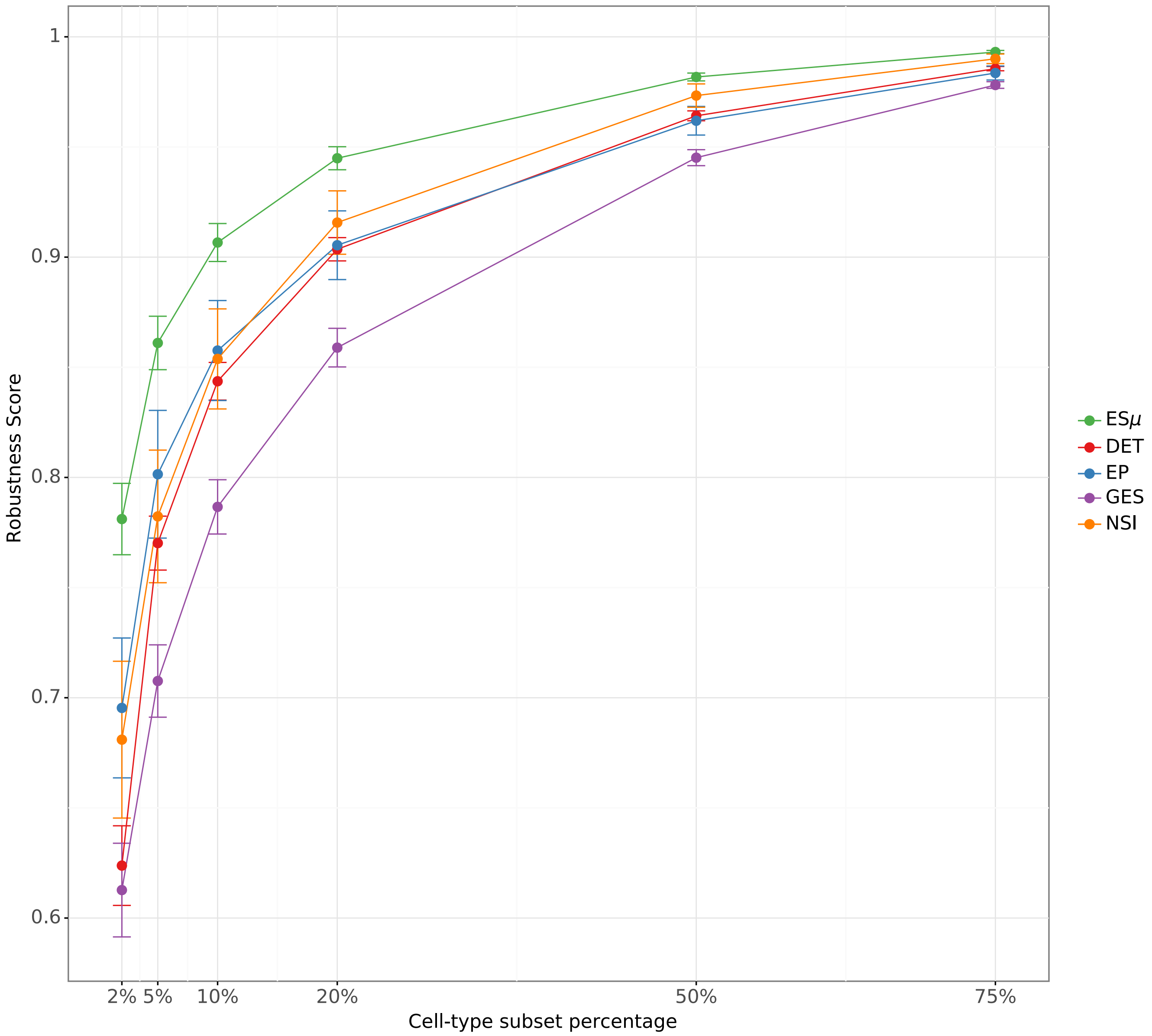

**Fig. S3.3 | Robustness of ES metrics.** The Fig. shows Robustness Score (RS) for each ES metric as a function of cell type subset percentage. Each point represents the mean RS across 12 focal cell types (see Methods). Error bars indicate standard error of the mean.

### Supplementary Note 4
### Cell type gene co-expression networks

#### Introduction
This Supplementary Note contains detailed methods related to cell type gene co-expression networks and their limitations.

#### Limitations of identifying co-expression networks from scRNA-seq data
In this section we discuss the relevance and limitations of identifying gene co-expression networks from cell type expression data. Some of these limitations have also been discussed elsewhere [40,41] and may contribute to the lack of identified BMI prioritized cell type gene co-expression networks.

While WGCNA has primarily been applied to bulk RNA-seq expression data, several recent studies have demonstrated its effectiveness in single-cell data [42,43]. As gene-gene correlations reflect gene expression variability, WGCNA analysis of heterogeneous bulk expression data consisting of several distinct cell type populations, will result in a coarse set of genes modules, largely reflecting cell type heterogeneity. WGCNA analysis of pure cell type populations from scRNA-seq data will result in more specific genes modules. However, constructing gene networks on pure cell type populations poses another problem: the true gene-gene correlations are more difficult to estimate in expression data with limited gene expression heterogeneity (expression variability). In addition, scRNA-seq data have increased technical noise (dropout effects) compared to bulk RNA-seq data, which reduces the ability to estimate the true gene-gene correlations.

### Supplementary Methods

#### Cell type labels

For the Mouse Nervous System dataset, we used Zeisel et al. (2018) cell type annotations: the first two letters in each cell type abbreviation denote the developmental compartment (ME, mesencephalon; DE, diencephalon; TE, telencephalon), letters three to five denote the neurotransmitter type (INH, inhibitory; GLU, glutamatergic) and the numerical suffix represents an arbitrary number assigned to the given cell type. Likewise, for the Tabular Muris dataset, we used the cell type labels as reported in their paper. For the six hypothalamic datasets, we added a label to more allow the reader to more easily understand, from which part of the hypothalamus a given cell type was sampled in the original study ("ARCME", arcuate nucleus median eminence complex; "HYPC", hypothalamus [44]; "HYPR", hypothalamus [45]; "LHA", lateral hypothalamus; "POA", preoptic area; "VMH", ventromedial nucleus) and the cell type it was annotated to in the original work (**Supplementary Table 12**).

#### Mouse Nervous System neurotransmitter annotation

We used the "Neurotransmitter" column of the cell type metadata (provided by Zeisel et al. (2018) accompanying mousebrain.org website) to group neuronal cell types into six neurotransmitter classes (transmitter listed in parenthesis): "excitatory" (glutamate), "inhibitory" (GABA or glycine), "monoamines" (adrenaline, noradrenaline, dopamine, serotonin), "acetylcholine" (acetylcholine), "nitric oxide" (nitric oxide) and "undefined" for neurons not matching these classes or without neurotransmitter data. When cell types were annotated with multiple transmitter classes in the "Neurotransmitter" column (e.g. glutamate and adrenaline), excitatory or inhibitory class took precedence in our assignment. See **Supplementary Fig. 3** for the neurotransmitter enrichment results.

#### Cell type gene co-expression networks

We identified cell type gene co-expression networks using a modified version of the robust WGCNA (rWGCNA) framework proposed by Gandal et al. (2018). To identify gene co-expression networks (henceforth referred to as *gene modules*) operating within a cell type, the input to rWGCNA was expression data for individual cell types. Our framework consists of the following steps:

**Pre-processing.** We used the *Seurat* R package (version 2.3). We filtered out genes expressed in fewer than 20 cells and removed any cell clusters containing fewer than 50 cells. The *NormalizeData()* function was used for normalizing raw expression values to a common transcript count (with n=10,000 transcripts as a scaling factor), log-transformation (log(x+1)), before scaling and centering with the *ScaleData()* command to arrive at a matrix of Z-scores. Principal component analysis was carried out using the *RunPCA()* function to find 120 Principal Components (PCs). Genes were then ranked by their highest absolute loading value on any given PC and the top 5,000 genes within each cell cluster were selected for co-expression analysis. We mapped mouse genes to orthologous human genes using Ensembl (v. 91), keeping only 1-1 mapping orthologs as done for the ES calcuations.

**Robust Weighted Gene Correlation Network Analysis and adjacency matrix calculation**. We used the *WGCNA* R package (version 1.66). For computing the gene-gene adjacency matrix we used the Pearson correlation coefficient and the *signed hybrid network* parameter. The *pickSoftThreshold()* command was used to identify soft thresholding powers. Powers corresponding to the top 95$^{th}$ percentile of network connectivity or above were discarded and the lowest soft threshold power between 1 and 30 to achieve a scale free topology R squared fit of 0.93 was selected; if no powers reached 0.93, the thresholding power with the highest R squared was chosen instead.

**rWGCNA consensus topological overlap matrix calculation.** The expression data was resampled as described in Gandal et al. (2018; drawing two thirds of the cells at random without replacement 100 times). The *consensusTOM()* function was used with a *consensusQuantile* of 0.2 to compute a signed consensus *topological overlap matrix* (TOM), and genes were then filtered using the output of the *goodGenesMS()* function called by *consensusTOM().*

**rWGCNA hierarchical clustering**. The consensus TOM matrices were converted to distance matrices and the *hclust()* function was used with the *average* method to cluster genes hierarchically. The *cutreeHybrid()* function was used with a *deepSplit* of 2, *minClusterSize* of 15 and *pamStage* set to TRUE to carve the dendrogram into modules. The *mergeCloseModules()* function was used to compute module eigengenes, the vector of cell embeddings on the first principal component of each module's expression submatrix. The same function was used to merge modules, using a *cutHeight* of 0.15 or less, corresponding to a Pearson correlation between module eigengenes of 0.85 or greater.

**rWGCNA gene-module connectivity** The module eigengenes and expression matrices were used with the *signedKME()* function to compute gene-module Pearson correlations, or *kMEs*, a measure of how close each gene is to each module. To ensure tightly connected modules, genes whose correlations with their assigned modules eigengene was not statistically significant after correcting for multiple testing, using the Benjamini-Hochberg false discovery rate (FDR) method, were removed.

### Supplementary Results

#### Expression specificity of known marker genes

First, to validate that our ES approach was able to delineate cell type-specific genes, we, for each of the four ES metrics, computed $ES_w$ estimates across four cell types with genes known to be specifically expressed in these cell types, namely hepatocytes (Apoa2), pancreatic alpha-cells (Gcg), striatum medium spiny neurons (Drd2) and mediobasal hypothalamic agouti related peptide (Agrp)-expressing neurons (Agrp). The four $ES_w$ metrics and the combined $ES_\mu$ metric correctly ranked the relevant genes at the top (Main text **Fig. 1d**). Conversely, plotting $ES_\mu$ values for these four genes across all cell types revealed that hepatocytes and alpha-cells exhibited the highest $ES_\mu$ for Apoa2 and Gcg, respectively, and that medium spiny neurons and Agrp-positive neurons exhibited the highest $ES_\mu$ for Drd2 and Agrp, respectively (Main text **Fig. 1e**).

#### TEINH12 enrichment

The TEINH12 cell type, a cholecystokinin (Cck)-expression interneuron, annotated to the hippocampus and cortex, clustered separately from the hippocampal and hypothalamus, thalamus and midbrain cell types suggesting partly overlapping transcriptional signatures with both brain regions. Interestingly, besides Cck the interneuron cell type exhibited specific expression of Ntrk2 (neurotrophic receptor tyrosine kinase 2), the Bdnf (brain-derived neurotrophic factor) receptor; two proteins encoded by genes with mutations that have been associated with obesity in humans and rodents [47] (**Supplementary Fig. 10 and Supplementary Table 6**). Previous work has suggested that Cck interneurons from the limbic system target the hypothalamus and knock-out of Ntrk2 in Cck-expressing interneurons has been linked to glucocorticoid resistance and mature-onset of obesity in mice [48].

#### Heritability of prioritized BMI cell types

In CELLECT we utilized a continuous representation of cell type expression, assuming there is a positive relationship between genes' ES values and their importance for a given trait. Stratifying genes into quintiles based on their ES values for a given cell type and calculating the enrichment of BMI heritability for each stratum, showed that an increase in ES was reflected in an increase in BMI heritability for the BMI-prioritized cell types (**Supplementary Fig. 11a; first row**). Other well-powered UKBB anthropometric traits GWAS did not exhibit any relationship between ES quintile and trait heritability for the BMI-prioritized cell types (**Supplementary Fig. 11a; second and third row**). These results indicate, that our models were well calibrated. To better understand the proportion of BMI heritability explained by variants mapped to a certain cell type, we compared the proportion of BMI heritability explained by each of the 10 cell types to the proportion trait-heritability explained by cell types known to play a key role in the given trait or disease. We found that the most enriched cell type for BMI, was the cortical TEGLU4 cell type, which comprised specifically expressed genes with genetic variants accounting for 28.5% of the SNP heritability (h2g) for BMI (**Supplementary Fig. 11b)**. Similarly, we estimated that SNPs mapped to genes specifically expressed in the pancreatic beta cells, an insulin secreting cell type playing a pivotal role in type 2 diabetes, explained 27.8% of the heritability for type 2 diabetes; hepatocytes explained 27.1% in the heritability for low-density lipoprotein [49]; T cells explained 20.7% of the heritability for rheumatoid arthritis [50]; and, finally, mesenchymal stem cells explained 20.4% of the heritability for human height [51] (**Supplementary Fig. 11b**). These results show that genetic susceptibility to obesity conferred by variants mapped to genes specifically expressed in the cortical cell type roughly corresponds to the heritability conferred by pancreatic beta cell variants on type 2 diabetes.

### Supplementary Figures

a

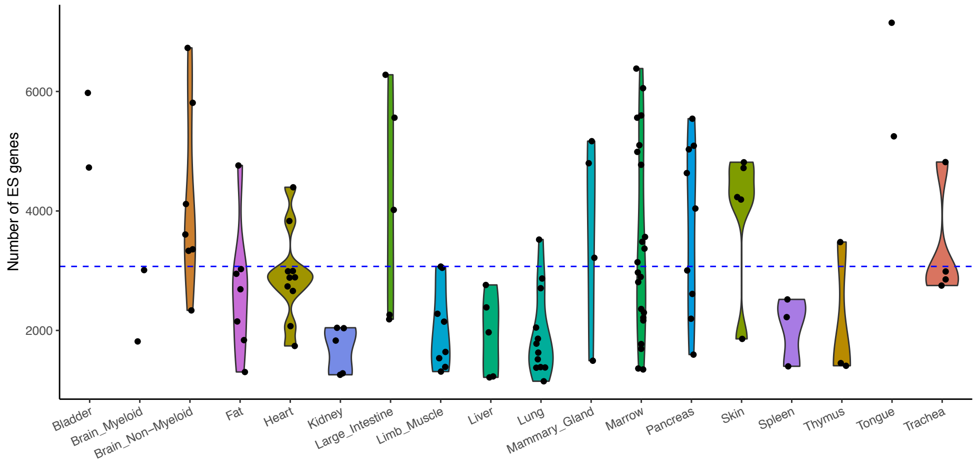

b

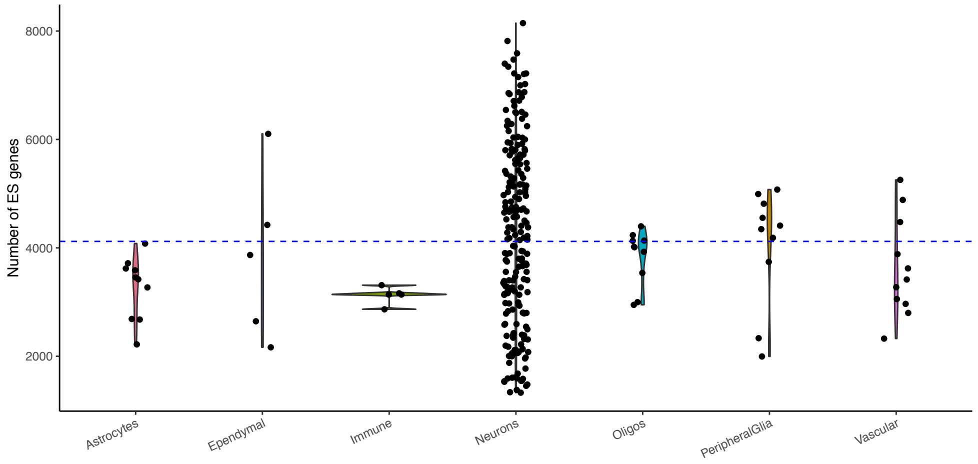

c

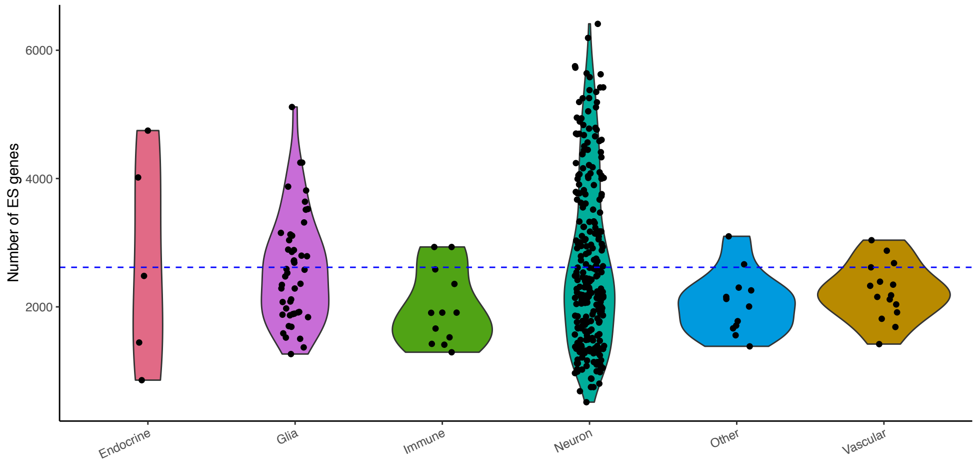

**Supplementary Fig. 1: Number of ES genes**

Distribution of the number of *ES genes* across cell type categories. Points represent cell types. The horizontal blue line reflects the global mean across all cell types. **a,** Tabula Muris (cell types grouped by tissue). **b,** Mouse Nervous System (cell types grouped by class). **c,** Hypothalamus datasets (cell types grouped by class).

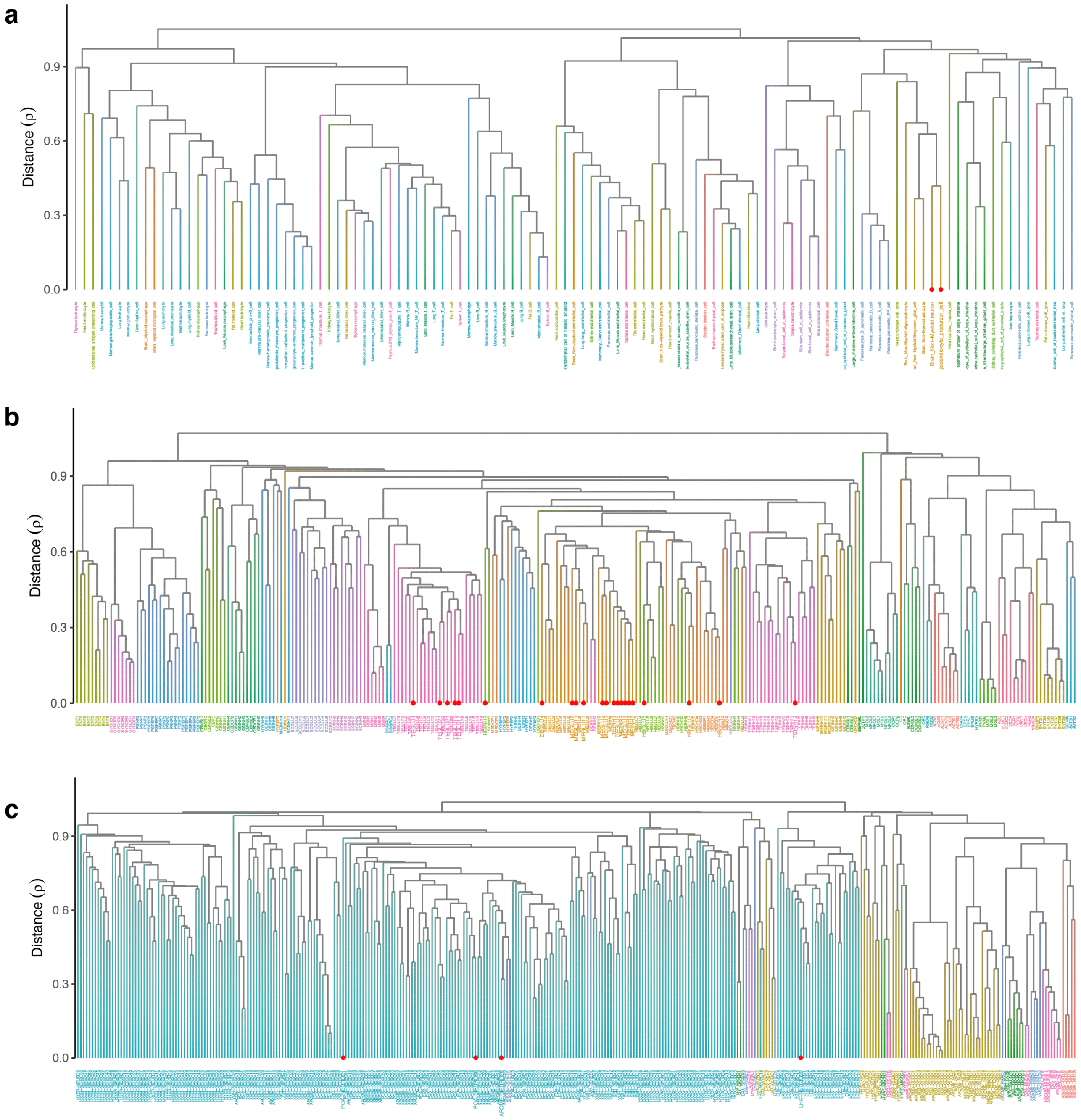

**Supplementary Fig. 2: Hierarchical clustering of cell types using $ES_{\mu}$**
Dendrogram of cell types clustered using average linkage of $ES_{\mu}$ Pearson's correlation. Dendrograms are shown for **a,** Tabula Muris; **b,** Mouse Nervous System; and **c,** Hypothalamus datasets. Cell types highlighted by red points (positioned at leaf nodes) passed the Bonferroni significance threshold in BMI GWAS prioritization analysis.

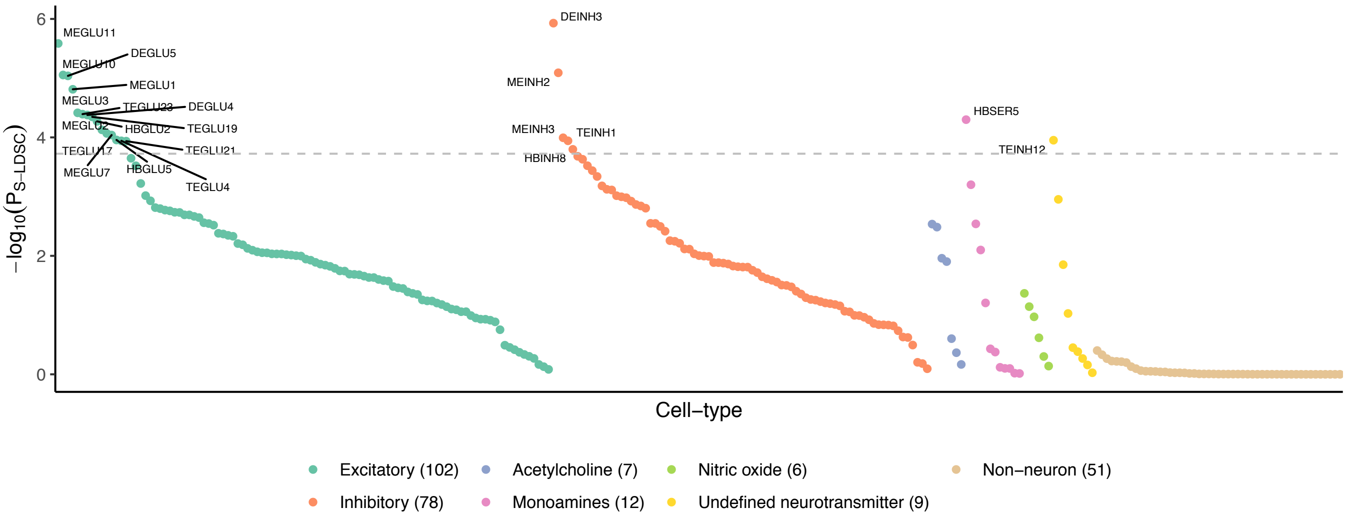

**Supplementary Fig. 3: BMI-prioritized Mouse Nervous System cell type neurotransmitter classes**

BMI GWAS prioritized cell types prioritization enriched for neurons. Non-neuronal (glial and vascular) cells did not exhibit any genetic enrichment. Cell types were grouped by neurotransmitter and ordered by genetic prioritization *P*-value ($P_{\text{S-LDSC}}$). See Supplementary Methods for a description of neurotransmitter annotation. The horizontal line marks the Bonferroni significance threshold ($P_{\text{S-LDSC}}<0.05/265$).

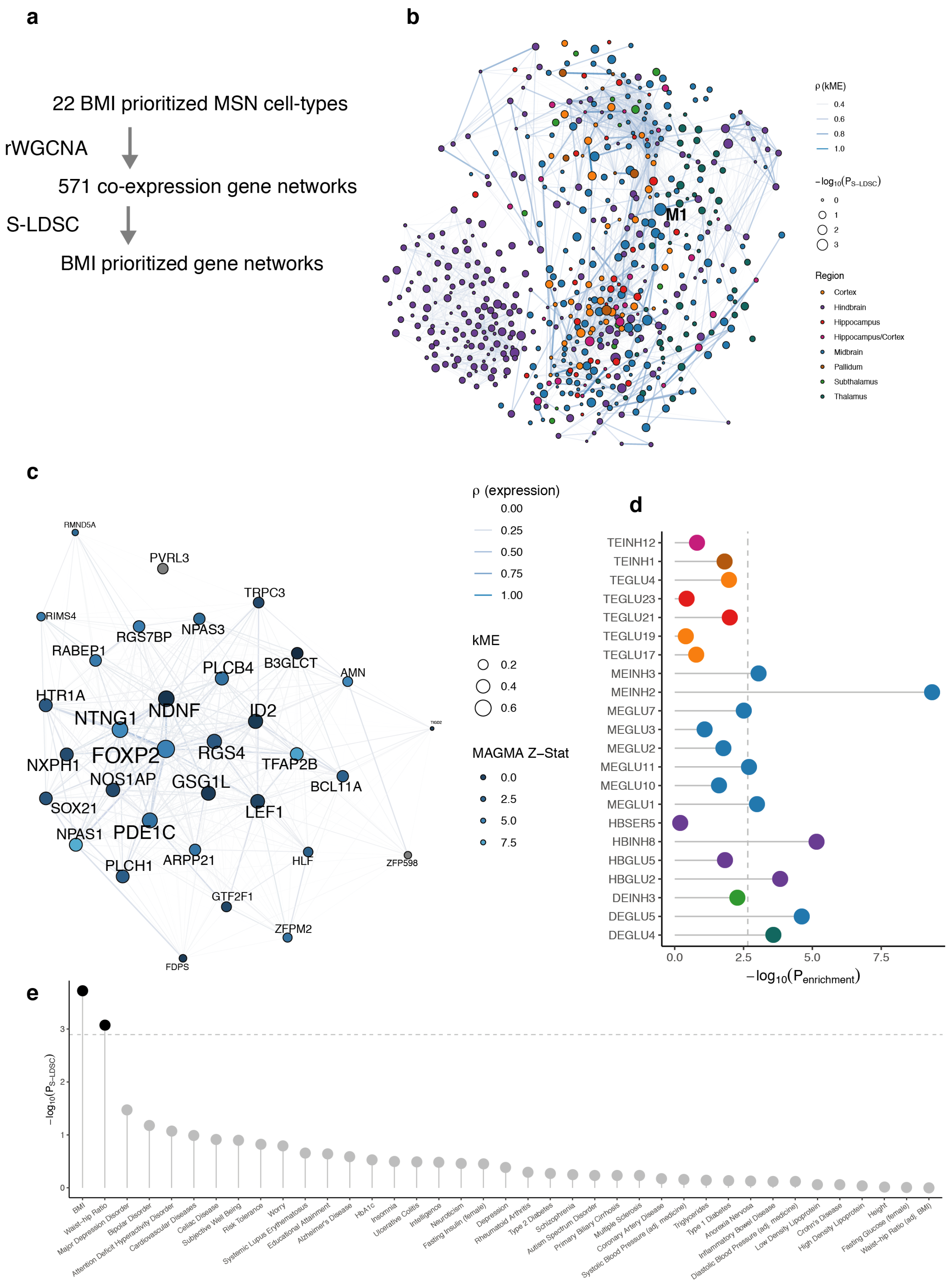

*See legend on the next page*

**Supplementary Fig. 4 (previous page): Genetic prioritization of cell type gene co-expression networks**

**a,** Overview of our approach to identify and prioritize cell type gene co-expression networks (modules). We used robust Weighted Gene Correlation Network Analysis (rWGCNA) to identify gene modules on expression data from individual BMI-prioritized cell types. The resulting modules were used as input to S-LDSC for genetic prioritization. **b,** Network visualization of the 571 cell type gene modules. The graph shows the absolute Pearson's correlation (ρ, edge width) between modules (nodes). Node color indicates the region of the cell type from which the module originates; node size represent genetic prioritization *P*-value for BMI (-$\log_{10}(P_{\text{S-LDSC}})$). The M1 module (originating from the MEINH2 cell type) is highlighted as the top significant module. The M1 module is not highly correlated with other modules. Only edges with ρ>0.3 are shown. **c,** Network visualization of the M1 gene module. The graph shows the absolute Pearson's correlation (ρ, edge width) between genes (nodes) in the module. Node size indicate kME value (a measure of gene-module membership); node color indicates MAGMA BMI gene Z-statistic (an aggregated measure of nearby variants BMI association). **d,** Enrichment of M1 module genes in BMI-prioritized cell types. Genes in the M1 module are enriched among expression specific genes for multiple prioritized cell types, but most strongly in MEINH2. The dashed line indicates the Bonferroni significance threshold ($P_{\text{enrichment}}<0.05/22$). **e,** Genetic prioritization of M1 module across multiple GWAS traits. The M1 module is associated with BMI and waist-hip ratio (passing Bonferroni significance threshold $P_{\text{S-LDSC}}<0.05/39$).

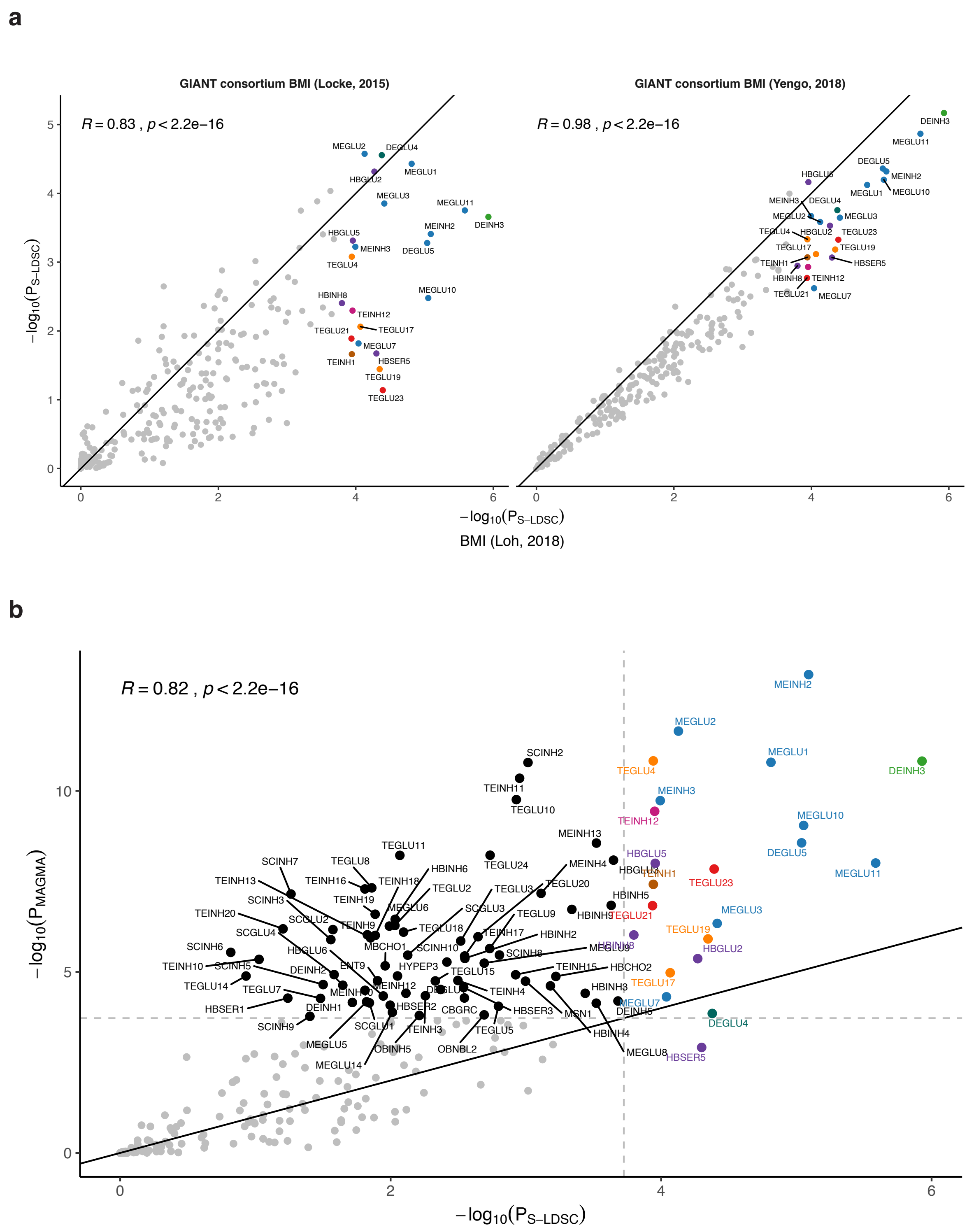

**Supplementary Fig. 5: Robustness of cell type prioritization results**

**a,** Comparison of Mouse Nervous System cell type prioritization results between the primary BMI analysis (Loh *et al.* (2018) GWAS summary statistics) and the Locke *et al.* (2015) GWAS summary statistics (sample size >320,000; left plot) and Yengo *et al.* (2018) BMI GWAS summary statistics (sample size >680,000; right plot). Pearson's correlation ($R$) is shown in the top left corner. Solid line shows x=y. **b,** Comparison of Mouse Nervous System cell type BMI prioritization results obtained using MAGMA (y-axis) and S-LDSC (x-axis). Pearson's correlation ($R$) is shown in the top left corner. Solid line show x=y. Dashed lines highlight cell types passing the Bonferroni significance threshold.

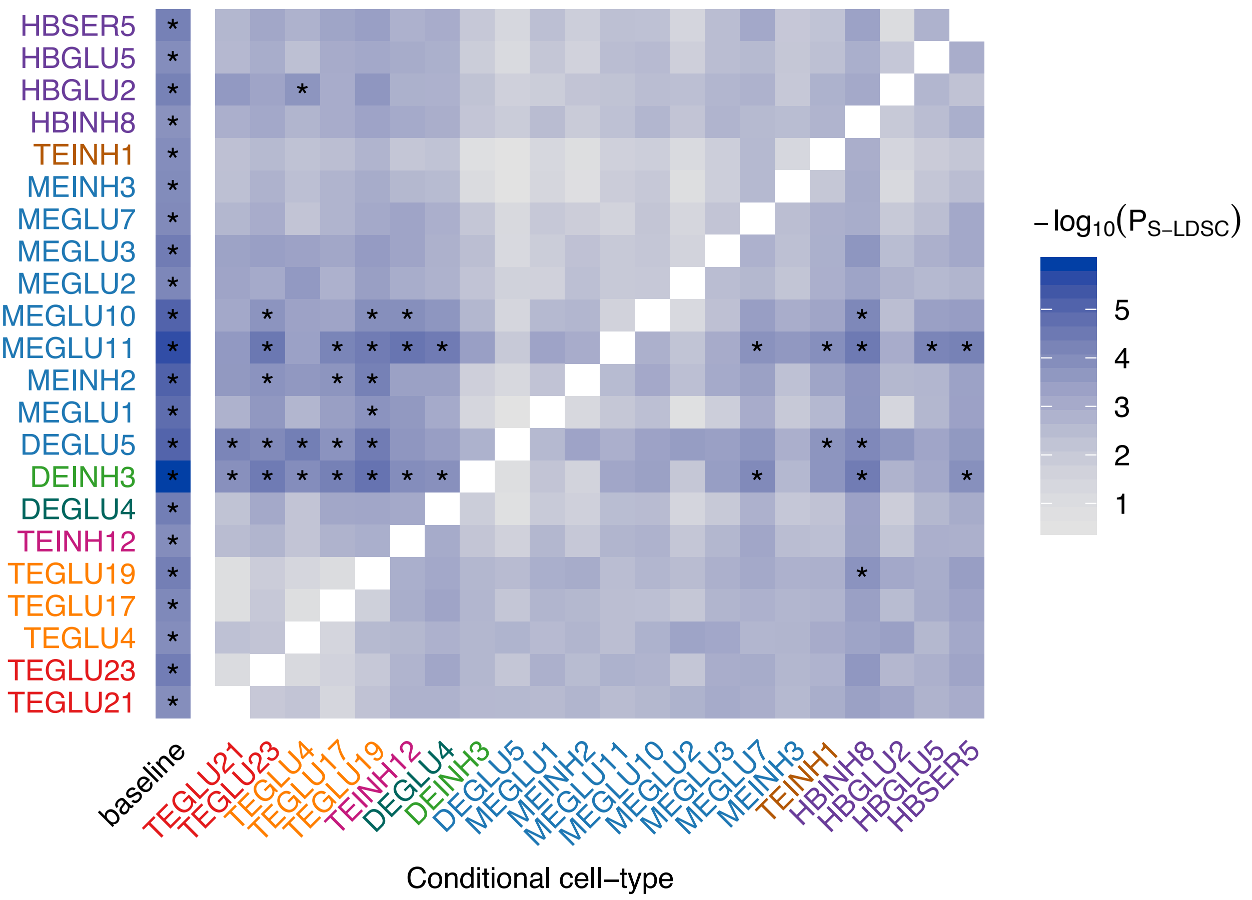

**Supplementary Fig. 6: Conditional analysis of BMI-prioritized Mouse Nervous System cell types**

Conditional genetic prioritization analysis of BMI-prioritized cell types. We used S-LDSC to re-estimate genetic prioritization *P*-values, conditioning on each BMI-prioritized cell type. Columns indicate the cell type conditioned on. The left column ('baseline') shows the unconditioned results (as shown in main text **Fig. 3a**). Cell-types are colored by their brain region as shown in main text **Fig. 3a**. NA values are colored in white (diagonal values) indicate cases were the prioritized and conditioned cell types are identical. Asterisks (*) mark *P*-values passing the Bonferroni significance threshold ($P_{S\text{-}LDSC}$<0.05/265) from main text **Fig. 3a**).

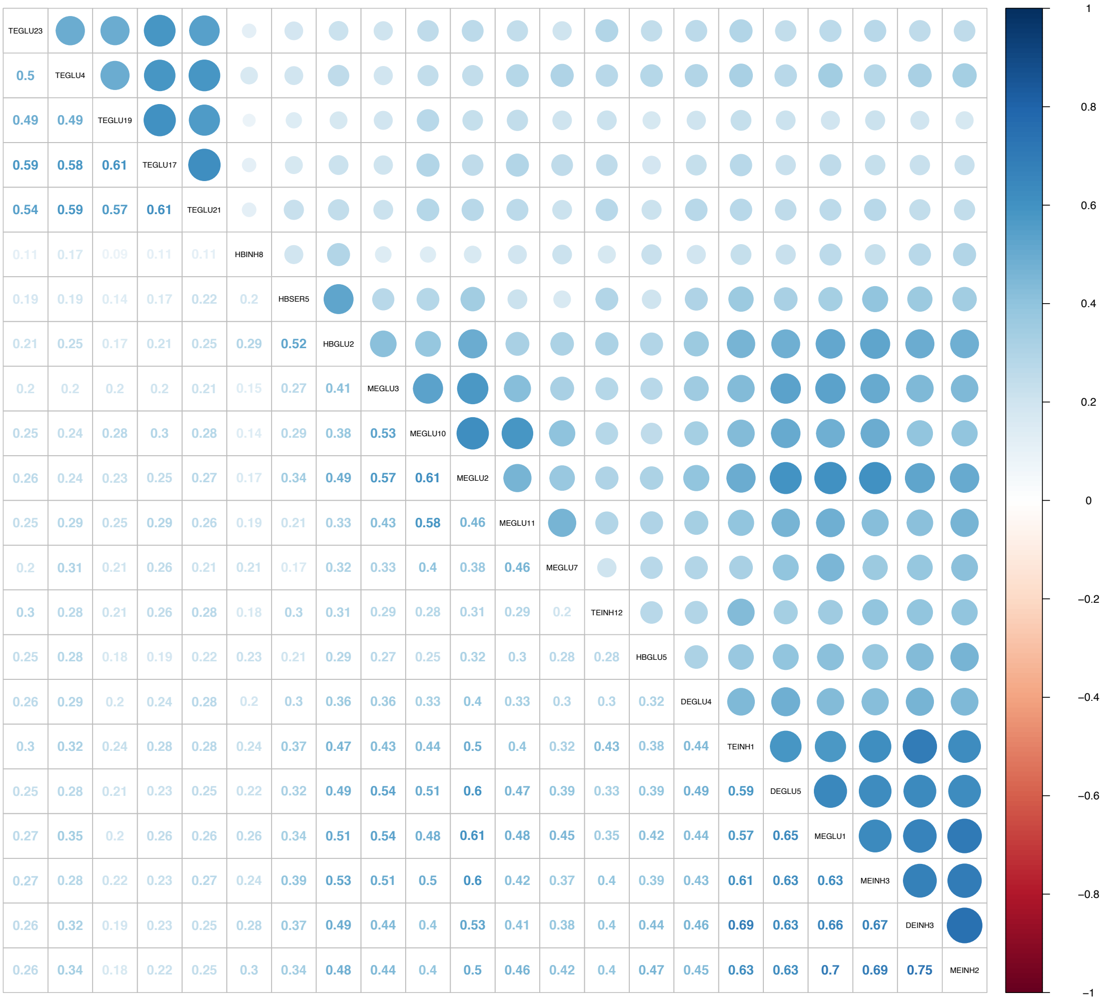

**Supplementary Fig. 7: Correlation of Mouse Nervous System BMI-prioritized cell types**
Correlogram of cell type $ES_\mu$ Pearson's correlations. Cell types are ordered by hierarchical clustered using Ward's method. The plot was generated using the "corrplot" R package.

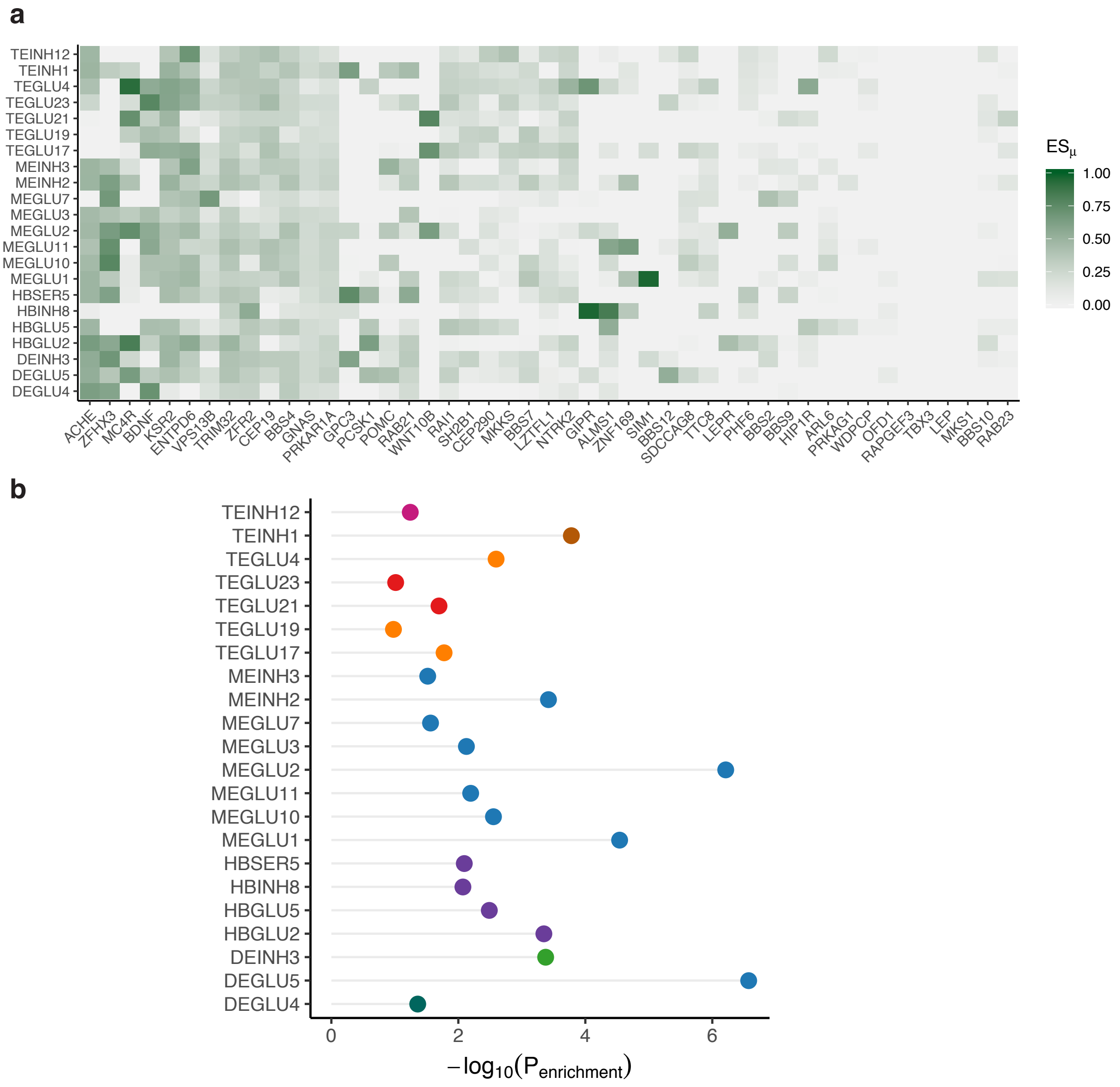

**Supplementary Fig. 8: Enrichment of rare variant obesity geneset for BMI-prioritized cell types**

**a,** Heatmap showing $ES_{\mu}$ values for genes in the obesity geneset (columns) for BMI-prioritized cell types (rows). **b,** Cell type enrichment of rare variant obesity geneset for BMI-prioritized cell types.

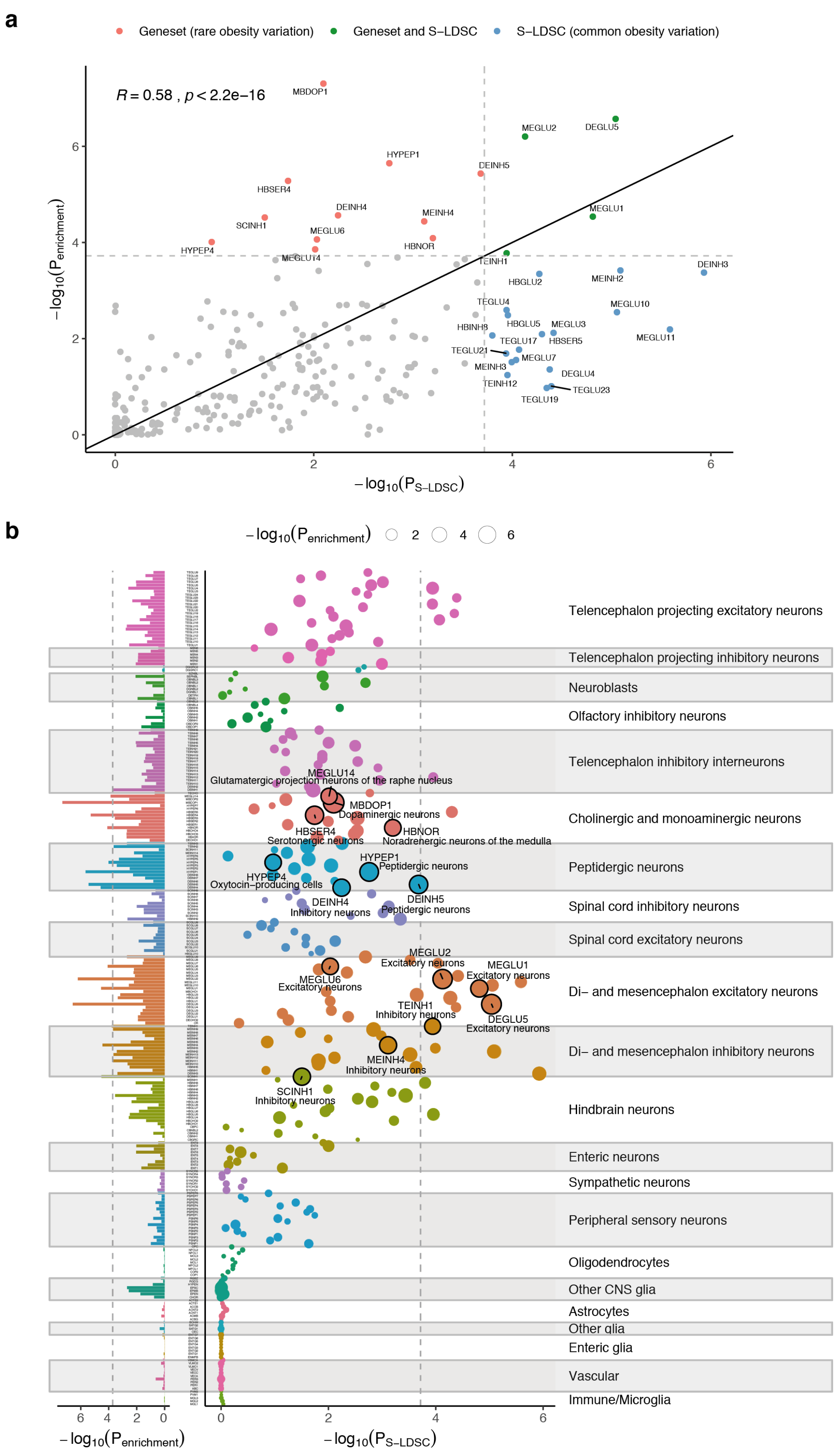

*See legend on the next page*

**Supplementary Fig. 9: Convergence of cell type prioritization based on common and rare variants**

**a,** Cell type prioritization of rare (bar chart, left part) and common variation (bubble chart, right part). The bar chart depict the cell type enrichment for the rare variant obesity geneset (genes predominantly derived from family studies of syndromic obesity and exome chip association analysis). The bubble chart depicts the BMI GWAS-based cell type genetic prioritization *P*-values (y-axis) as assessed using S-LDSC using the common BMI variants (MAF > 5%, identical to **Fig. 3a**; see Methods). Circle sizes represent the $-\log_{10}(P_{\text{enrichment}})$ cell type enrichment of rare variant obesity geneset. Circles with black edges mark cell types passing the Bonferroni significance threshold ($P_{\text{enrichment}}<0.05/265$). Bar chart (left plot) shows the distribution of $-\log_{10}(P_{\text{enrichment}})$ across all cell types. Dashed lines indicate Bonferroni significance threshold. Cell types are grouped by the cell type taxonomy shown in **Fig. 3b**. **b,** Comparison of cell type BMI GWAS-based prioritization and rare variant obesity geneset enrichment-based results. Cell type BMI GWAS-based prioritization (x-axis) and cell type enrichment of rare variant obesity genes (y-axis). Pearson's correlation (*R*) is shown in the top left corner. Dashed lines indicate cell types passing the Bonferroni significance threshold. Solid line shows x=y relationship.

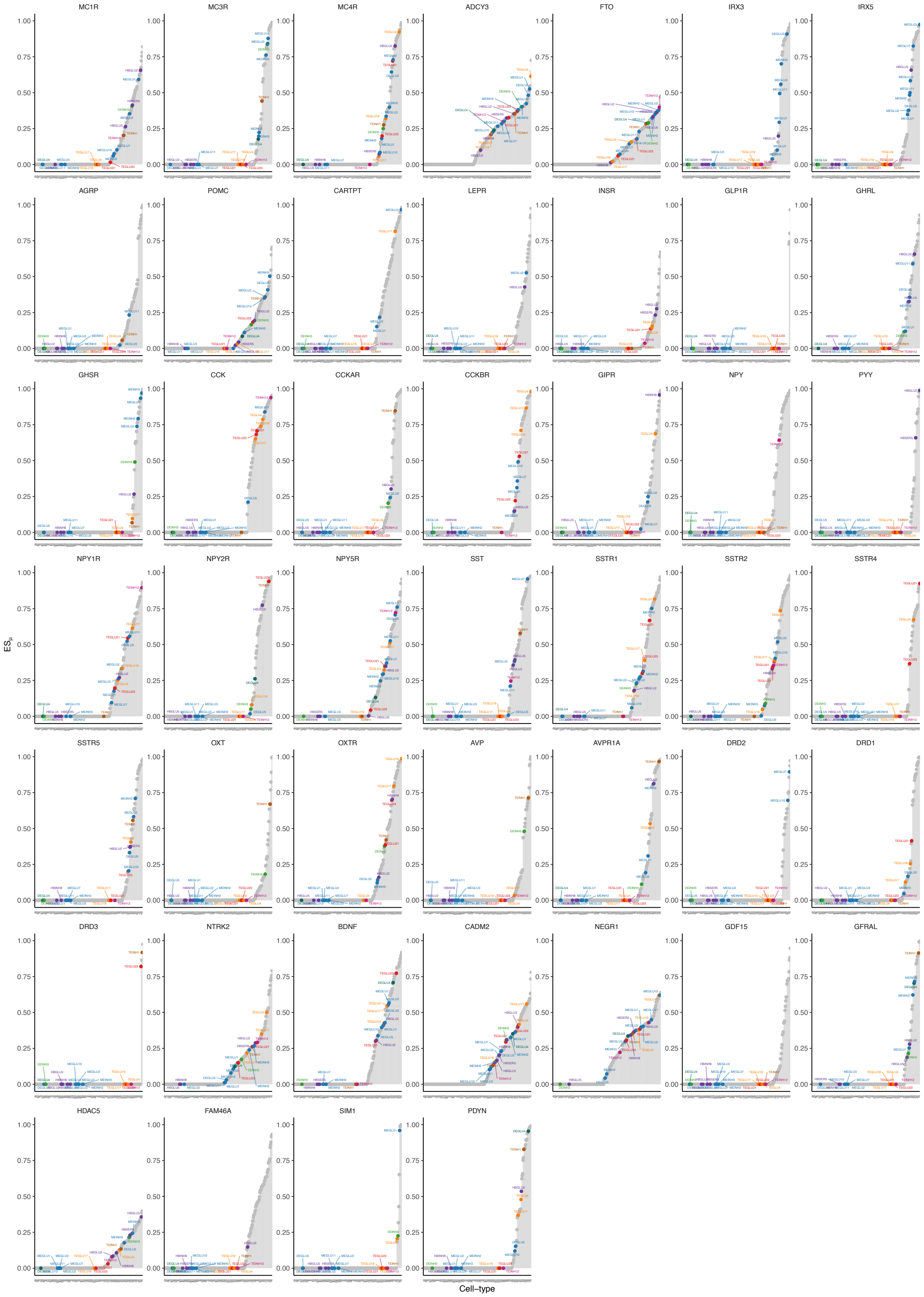

**Supplementary Fig. 10: $ES_\mu$ plot for selected genes**

$ES_\mu$ plots for genes selected based on their suggested role in appetite regulation, energy homeostasis or obesity. The plot shows gene $ES_\mu$ (y-axis) for each cell type (x-axis, ordered by increasing values of $ES_\mu$) and highlights the BMI-prioritized cell types.

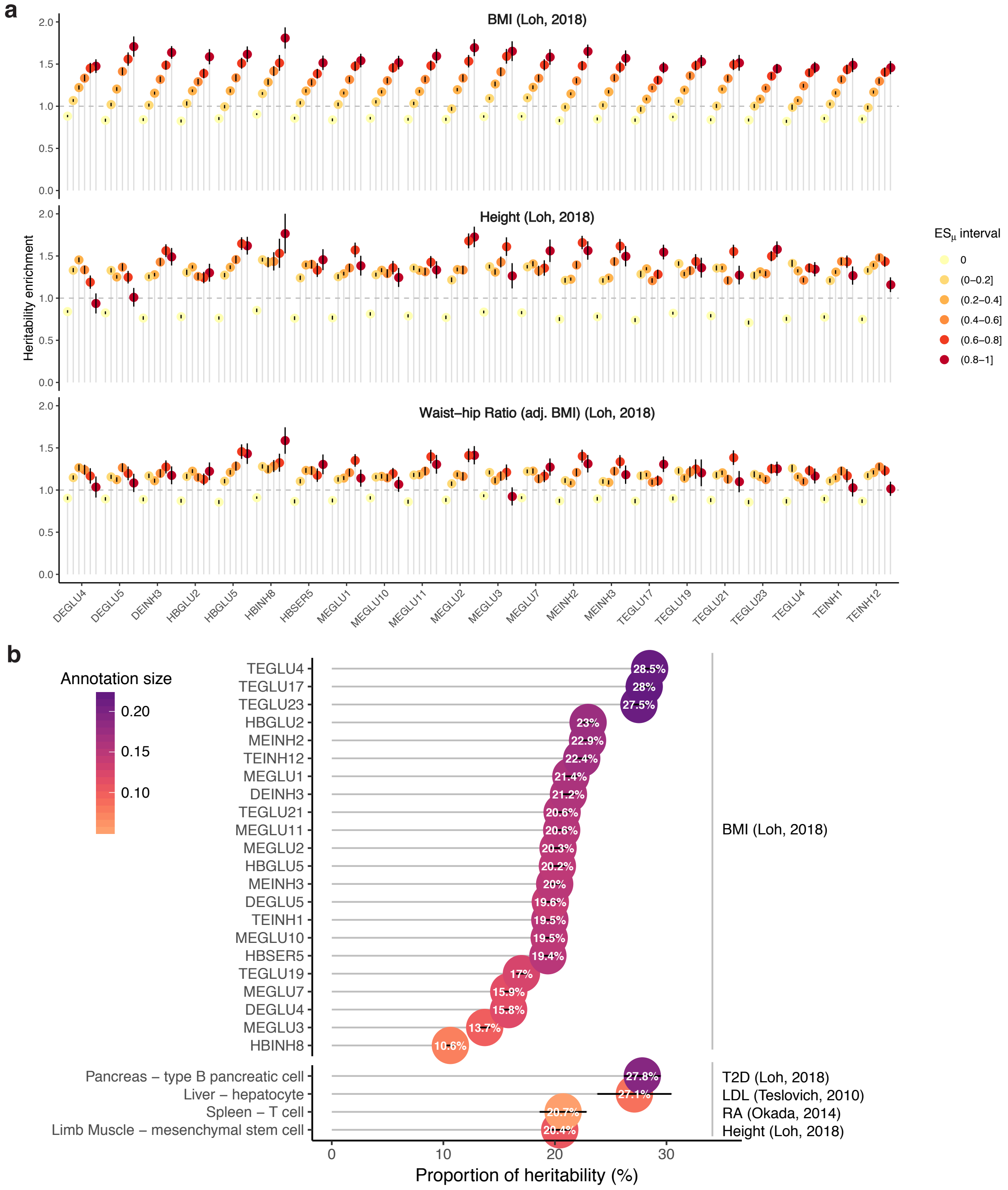

**Supplementary Fig. 11: Heritability of BMI prioritized cell types**

**a,** Heritability enrichment of cell type ESμ intervals. Heritability enrichment was estimated using S-LDSC on cell type $ES_\mu$ annotations partitioned into five equally spaced intervals and an interval including $ES_\mu$=0. The intervals 0 and (0.8-1] represent the heritability enrichment of by the variants with the lowest and highest $ES_\mu$ values, respectively. Error bars represent 95% confidence intervals. The top, middle and bottom panel show results for BMI, height and waist-hip ratio, respectively. BMI heritability enrichment increases with increasing $ES_\mu$ value for prioritized cell types. **b,** Proportion of BMI heritability explained by prioritized cell types. We used S-LDSC to estimate the proportion of trait SNP heritability explained by each cell type annotation. For comparison we report the proportion of heritability explained by cell types with known etiology for selected traits: type 2 diabetes (T2D), low-density lipoprotein (LDL), rheumatoid arthritis (RA) and human height. Circles are colored by annotation size reflecting the proportion of variants covered by the cell type annotation (a value of one means that all variants were covered). Error bars represent 95% confidence intervals.
